# Intergenerational transmission of longevity is not affected by other familial factors: *Evidence from 16,905 Dutch families from Zeeland, 1812-1962*

**DOI:** 10.1101/781500

**Authors:** R.J. Mourits, N.M.A. van den Berg, M. Rodríguez-Girondo, K. Mandemakers, P.E. Slagboom, M. Beekman, A.A.P.O. Janssens

## Abstract

Studies have shown that long-lived individuals seem to pass their survival advantage on to their offspring. Offspring of long-lived parents had a lifelong survival advantage over individuals without long-lived parents, making them more likely to become long-lived themselves. We test whether the survival advantage enjoyed by offspring of long-lived individuals is explained by environmental factors. 101,577 individuals from 16,905 families in the 1812-1886 Zeeland cohort were followed over time. To prevent that certain families were overrepresented in our data, disjoint family trees were selected. Offspring was included if the age at death of both parents was known. Our analyses show that multiple familial resources are associated with survival within the first 5 years of life, with stronger maternal than paternal effects. However, between ages 5 and 100 both parents contribute equally to offspring’s survival chances. After age 5, offspring of long-lived fathers and long-lived mothers had a 16-19% lower chance of dying at any given point in time than individuals without long-lived parents. This survival advantage is most likely genetic in nature, as it could not be explained by other, tested familial resources and is transmitted equally by fathers and mothers.

## 1. Introduction

Living a long and healthy life is a dream many share. Yet, some of us live longer than others and reach advanced ages in better health. To the longest-lived individuals in this world age-related diseases – such as diabetes, cardiovascular diseases, Alzheimer, etc. – seem to be a less heavy burden. On top of that, studies have shown that long-lived individuals seem to pass their survival advantage on to their offspring (Atzmon et al., 2004; Christensen et al., 2008; Dutta et al., 2014; Gjonça & Zaninotto, 2008; Newman et al., 2011; Terry et al., 2004; Terry et al., 2008; Van den Berg et al., 2018; Westendorp et al., 2009). Offspring of long-lived parents has a lifelong survival advantage over individuals without long-lived parents, making them more likely to become long-lived themselves (Dutta et al., 2013; Gudmundsson et al., 2000; Houde, Tremblay & Vézina, 2008; Perls et al., 2002; Terry et al., 2004; Van den Berg et al., 2018; Westendorp et al., 2009, Willcox et al., 2006). It is widely believed that members of these families have a genetic predisposition that benefits their own as well as their offspring’s survival (Sebastiani et al., 2015; Shadyab & LaCroix, 2015; Van den Berg et al., 2018). Yet, there are also other familial factors that have been associated with longevity, e.g. parity, farming, social class, or smoking and drinking behavior (Gavrilov & Gavrilova, 2015; Kerber et al., 2001; Robine et al., 2003; Sun et al., 2015; Tabatabaie et al., 2011; Temby & Smith, 2014). These factors are often shared between parents and children and, as such, can correlate a family’s chances to become long-lived (Cournil, Legay & Schächter, 2000; Gavrilov & Gavrilova, 2015; Matthijs, Van de Putte & Vlietinck, 2002; Montesanto et al., 2017; Temby & Smith, 2014; You, Gu & Yi, 2010). Currently, studies have found little evidence that social factors affected the association between parental longevity and offspring survival (Gavrilov & Gavrilova, 2015; You, Gu & Yi, 2010). However, these studies were based on very specific populations, relatively small samples, and had limited information on familial resources. Therefore, to what extent other familial factors affect the association between parental longevity and offspring survival is open for discussion.

Familial resources play an important role in determining their offspring’s survival. Multiple familial factors act in utero and during the early stages of life when offspring is very sensitive and dependent on their parents for survival (Ben-shlomo & Kuh, 2002; Elo & Preston, 1992; Smith & Hanson, 2015). Hence, losing a parent or high mortality among siblings early in life are known to negatively affect survival, whereas having a healthy mother or growing up in the right socioeconomic environment might benefit survival (e.g. Barker, 1990; Elo & Preston, 1992). Moreover, individual factors that have been associated with longevity – such as parity (Tabatabaie et al., 2011; Westendorp & Kirkwood, 1998), age at last birth (Sun et al., 2015), social class (Gavrilov & Gavrilova, 2015; Temby & Smith, 2014), smoking, and drinking (Kerber et al., 2001; Sun et al., 2015; Temby & Smith, 2014) – are passed on from parents to children. Just like parental longevity, these resources can cluster within families and are transferred to future generations (Broström, Edvinsson & Engberg, 2018; Knigge, 2016; Morris et al., 2011; Sommerseth, 2018; Van Dijk & Mandemakers, 2018). Accordingly, the survival advantage of having a long-lived parent can be caused by inherited genetic predispositions or environmental conditions due to high infant mortality in the sibship, familial fertility histories, or shared socioeconomic resources. This begs the question whether the intergenerational transmission of longevity is affected by any of these familial factors. And, if so, which of the familial resources is most important.

In this paper we explore whether high infant mortality in the sibship, familial fertility histories, and shared socioeconomic resources affect the association between parental longevity and offspring survival. We use reconstituted family data from the historical dataset LINKS-Zeeland (Mandemakers & Laan, 2017) to study 16,905 disjoint families with 101,577 children. These children were born between 1812 and 1886 and lived and died in the Dutch province of Zeeland, which – at the time – was known for its high fertility and high infant mortality. Within this context, we first verify the known relation between parental longevity and offspring survival. We group offspring by their parent’s longevity and show survival plots for offspring with no parents belonging to the top 10% survivors, a top 10% surviving father, a top 10% surviving mother, and two top 10% surviving parents. Second, we enquire whether the survival advantage enjoyed by offspring of long-lived parents is dependent on high infant mortality in the sibship, familial fertility histories, and shared socioeconomic resources. Third, we determine which familial resources are most important for the survival of offspring and show that the association between familial resources and offspring survival is remarkably stable over the life course. Finally, we discuss what these outcomes mean for research on longevity and historical demography.

## 2. Literature discussion

We already established that parental longevity is an important predictor of offspring survival. But besides parental longevity, high infant mortality in the sibship, familial fertility conditions, and shared socioeconomic resources are familial factors that might also associate with individual chances to become long-lived. High early-life mortality within the family can be an indication of familial frailty, an unhealthy living environment, or a mixture of both – e.g. increased vulnerability to environmental factors, such as epidemics or polluted drinking water – (Bengtsson & Lindström, 2000, 2003; Quaranta, 2013; Van Dijk, Janssens & Smith, 2018; Vaupel, 1988). A family’s fertility history gives insight into parental physical fitness at conception (Barker, 1990; Floud et al., 2011; Perls, Alpert & Fretts, 1997; Westendorp & Kirkwood, 1998; Wrigley, 2004). Socioeconomic status and family compositions indicate which resources are available to each household member (Bengtsson & van Poppel, 2011; Blake, 1981; Yerushalmy, 1938). In this section, we discuss how these three types of parental resources associate with offspring survival.

### 2.1 High infant mortality in the sibship

Offspring of long-lived parents is thought to obtain their parents’ predisposition towards longevity. Reversely, offspring that is confronted with high mortality in the family may have an inherited survival disadvantage. Having parents with a short lifespan can indicate that offspring had frail or genetically burdened parents (Vaupel, 1988). These parents with a short lifespan may have suffered from early onsets of degenerative diseases, which, if passed on, in turn reduce survival of offspring. Furthermore, poor maternal health can also negatively affect the in-utero development of offspring, resulting in survival disadvantages later in life (see e.g. Barker, 1990; Rosano, Botto, Botting, Mastroiacovo & Germany, 2000; Smith & Hanson, 2015). Offspring is also directly affected by the death of a parent. The loss of one’s parents meant the loss of food, care, and possibly future chances on the marriage and labor market (Cooper, 1992, p. 296; Kok & Delger, 1998; Van Poppel, de Jong & Liefbroer, 1998). Especially the loss of one’s mother at a young age was detrimental and could result in an early demise (Rosenbaum-Feldbrügge, 2018). Moreover, losing a parent at a young age seems to speed up reproduction and has been thought to increase mortality levels as well. Multiple studies have shown that the early loss of a parent introduces earlier puberty (Bogaert, 2008; Webster et al., 2014) and a tendency to reproduce earlier (Störmer & Lummaa, 2014; Voland & Willführ, 2017). However, studies on the effect of losing a parent on later-life mortality outcomes have shown mixed results (Campbell & Lee, 2009; Gagnon & Mazan, 2009; Smith et al., 2009b; 2014; Todd, Valleron & Bougnères, 2017; Van Poppel, De Jong & Liefbroer, 1998; Willführ, 2009). Thus, it is uncertain whether the early loss of a parent affected offspring survival after the initial shock.

Besides having a short-lived parent, having multiple siblings who died in infancy can also be a sign of inherited frailty. Increased risks on infant and child mortality with “socioeconomic, genetic, behavioral, and environmental roots” (Van Dijk, 2018) are passed on to future generations and can shorten the lives of offspring as well as grandchildren (Broström, Edvinsson & Engberg, 2018; Gagnon et al., 2009; Hin, Ogórek & Hedefalk, 2016; Janssens, Messelink & Need, 2010; Sommerseth, 2018; Van Dijk & Mandemakers, 2018). To a large extent high infant mortality was determined by the living environment. Infant mortality differed widely between and within countries and ranged from less than 5% to around 40% of all newborns (Klüsener et al., 2014; van den Boomen & Ekamper, 2015; Van Poppel, Jonker & Mandemakers, 2005). Growing up in an unhealthy environment not only determined how many infants died, but also scarred the survivors. Even years after being exposed to outbreaks of infectious diseases, survivors showed increased mortality risks (Bengtsson & Lindström, 2003; Quaranta, 2013). Both in high and low mortality environments, mortality rates were significantly higher for the survivors from high mortality families (Van Dijk, Janssens & Smith, 2018). Therefore, high levels of infant mortality in the family should be seen as an indicator of inherited frailty as well as an unhealthy living environment.

### 2.2 Familial fertility history

Besides infant mortality in the sibship, familial fertility histories might play an important role in determining the survival of offspring. Offspring survival is linked to parental fertility through reproductive ageing, genetic predispositions, and parental physical fitness. Reproductive ageing stresses the benefits of having a mother who is able to reproduce until advanced ages. Under natural fertility conditions, most women give birth to their last child between ages 35 and 45, while last births after age 45 are rare (Eijkemans et al., 2014). Late-reproducing women are not only able to give birth at advanced ages, but also show lower mortality rates after age 50 than other women and are more likely to become centenarians (Gagnon et al., 2009; Helle, Lummaa & Jokela, 2005; Smith et al., 2009a; Smith, Mineau & Bean, 2002; Sun et al., 2015). The causal mechanism that links age at last birth and survival after age 50 has yet to be established (Gagnon, 2015), but is thought to be rooted in social and economic benefits or beneficial genetic predispositions (Te Velde & Pearson, 2002). A mother who was able to conceive children at advanced ages can pass these characteristics on to her offspring, as both social position and female ages at last birth/menopause cluster within families (Knigge, 2016; Zimmer, Hanson & Smith, 2016; De Bruin et al., 2001; Morris et al. 2011; Pettay et al., 2005; Van Asselt et al., 2004; Walter et al., 2012). Furthermore, late-reproducing mothers might have been healthier and were able to give birth to babies with a higher birth weight, which would make her offspring less vulnerable and more resilient to all kinds of infectious diseases. Therefore, having a mother who was able to conceive children at advanced ages might be beneficial for her offspring’s survival.

Although having a mother who reproduced until advanced ages can be considered beneficial for her offspring, having a mother who has a large number of births might not be so beneficial for her offspring. The link between number of offspring and age at death itself is weak at best (Helle, Lummaa & Jokela, 2005; Hurt, Ronsmans & Thomas, 2006; Le Bourg, 2007) and seems to be strongly dependent on parental health and mortality during childbearing ages (Doblhammer & Oeppen, 2003). However, number of offspring is associated with short birth intervals and parental ageing, which are known to affect offspring survival (Dewey & Cohen, 2007; Gavrilov & Gavrilova, 2000; Kozuki et al., 2013; Xie et al., 2018). Giving multiple births in rapid succession can deplete a woman, due to increased exposure to stress, additional energy requirements, and having less time to recover (Engelen & Wolf, 2011; Winkvist, Rasmussen & Habicht, 1992). Gestation and childbirth take their toll on the female body, which can only recover with time and adequate nourishment. Shorter birth intervals indicate that women have less time to recover from a previous pregnancy, which is especially detrimental when access to food is restricted. This weakens a woman’s physiology and gives her an increased risk of having a miscarriage or giving birth to offspring with low birth weight. Therefore, having many children in a rapid succession will be detrimental to the mother’s and her offspring’s health, regardless of a possible genetic tradeoff between reproduction and lifespan (see e.g. Kirkwood, 1977; Westendorp & Kirkwood, 1998). Besides short birth intervals, number of offspring is also correlated with parental age at birth. Parental physical fitness decreases over time. Genetic damage accumulates over time and DNA mutations increase as parents grow older (Crow, 1993). Hence, older mothers’ germ cells contain more accumulated damage, whereas older fathers transmit germ cells that are more mutated. Due to the high mutation load, both can be detrimental to the health of their children and shorten their lifespan (Gavrilov & Gavrilova, 2000; Xie et al., 2018). Possibly as a result, later born children seem to show higher mortality rates (Barclay & Kolk, 2015; Engelen & Wolf, 2011; Hin, Ogórek & Hedefalk, 2016; Smith et al., 2014; Sommerseth, 2018; Van Dijk & Mandemakers, 2018).

Having parents who stopped reproducing early or who had long birth intervals are not necessarily signs of having a healthy family. In natural fertility populations, a low number of children is generally seen as an indicator of problems with parental fertility or parental health (Doblhammer & Oeppen, 2003). Weakened mothers were more likely to produce children with lower births weights and could pass on their frail physiology to their offspring. Furthermore, fertility problems can stem from genetic mutations in the parents’ germ cells that also negatively affect surviving offspring. Hence, offspring belonging to a small sibling set might be the result of inherited frailty or development problems. Conversely, however, having a healthy mother can instill offspring with a higher birthweight and concomitant survival advantages. It has been found that long-lived women age healthier than their non-long-lived counterparts (Atzmon et al., 2004; Christensen et al., 2008; Dutta et al., 2014; Gjonça & Zaninotto, 2008; Newman et al., 2011; Terry et al., 2004; 2008; Westendorp et al., 2009). Hence, women who have the potential to become long-lived might be better equipped to give birth to larger and healthier children who are less vulnerable to a wide range of environmental effects (Floud et al., 2011; Van den Berg et al., 2018; Wrigley, 2004). Following this line of reasoning, having a migrant mother could also increase offspring survival. Migrants are known to have a survival advantage compared to the general population, because they are healthier than the general population (Khlat & Courbage, 1996; Markides & Coreil, 1986; Puschmann, Donrovich & Matthijs, 2017; Wallace & Kulu, 2014). Hence, having a migrant mother might also increase her offspring’s chances to live a long life.

### 2.3 Shared socioeconomic resources

Parents determine the environments in which their children grow up. This is even more so in historical populations where parental socioeconomic status determined the quality, quantity and security of food as well as housing conditions and routine aspects of daily life. Daily nutrition is often thought to be one of the most important determinants to prevent and survive infectious diseases, which were more prevalent and virulent in the 19^th^ century (Fogel & Costa, 1997; McKeown, 1976; Preston, 1976; Rotberg & Rabb, 1985). Differences in access to food were considerable and caused differences in human stature: elite and farmer children were on average taller than their fellow countrymen, whereas children of laborers were shorter than the rest of the population (Alter & Oris, 2008; Beekink & Kok, 2017; Komlos, 1990; Mazzoni et al., 2017; Öberg, 2014; Ramon-Muñoz & Ramon-Muñoz, 2017). The relationship between parental socioeconomic status and mortality in the first five years of life seems to follow a similar pattern. Farmers’ and upper class children had a survival advantage over children from other parents (Breschi et al., 2011; Edvinsson et al., 2005; Janssens & Pelzer, 2012; Schumacher & Oris, 2011; Van Poppel, Jonker & Mandemakers, 2005). There is some evidence that the differences in survival between offspring of farmers, the elite, the middle class, and laborers remains present over the rest of the life course (Breschi et al., 2011; Hin, Ogórek & Hedefalk, 2016; Schenk & van Poppel, 2011; Van Den Berg, Lindeboom & Portrait, 2006). However, studies focused on the association between individual socioeconomic status and later-life survival in commercial-agricultural societies generally did not find any socioeconomic effects on differences in mortality rates (Bengtsson & van Poppel, 2011; Edvinsson & Broström, 2012).

Whether offspring was able to profit from parental resources is dependent on how resources were allocated in the household (Riswick, 2018). Children compete for their parent’s attention and resources in the household. Having multiple older brothers seems to be detrimental to survival in later life (Donrovich, Puschmann & Matthijs, 2014). Being born earlier in the birth order puts offspring in an advantageous position in terms of resources, as they have fewer siblings to compete with over available resources. Moreover, firstborn sons are much more likely to get paternal attention, as they are often supposed to take over their father’s trade and family’s assets, e.g. the family farm, smithy, bakery, or store. Earlier born siblings are known to have lower mortality rates (Barclay & Kolk, 2015; Engelen & Wolf, 2011; Hin, Ogórek & Hedefalk, 2016; Smith et al., 2014; Sommerseth, 2018; Van Dijk & Mandemakers, 2018), but it is not known whether this is caused by maternal depletion, resource competition, or selective parental investment.

### 2.4 Synthesis

In this paper, we test the discussed familial effects on offspring survival in one overarching framework. The effects of high infant mortality in the sibship, familial fertility histories, and shared socioeconomic resources are summarized in Figure 1. In each column we show the discussed mechanisms and their demographic indicators. In the upcoming paragraphs we test whether the association between parental longevity and offspring survival – defined as age at death – is affected by any of these factors. Besides effects of mortality, fertility, and socioeconomic resources, Figure 1 further shows possible associations between parental behavior and offspring survival. Parental behavior is not included in our analyses due to data constraints, but is known to be an indicator of offspring survival. Having a drinking or smoking parent seems harmful for offspring survival (Hill et al., 2000; Huizink & Mulder, 2006; Ji et al., 1997; Lindahl-Jacobsen et al., 2013), whereas breastfeeding might have been benefited offspring survival (van den Boomen & Ekamper, 2015; Walhout, 2010). During the period under observation, few people smoked, alcohol consumption was common practice, and breastfeeding practices varied considerably by region (Janssen & Van Poppel, 2015; van den Boomen & Ekamper, 2015).

**Figure 1:**
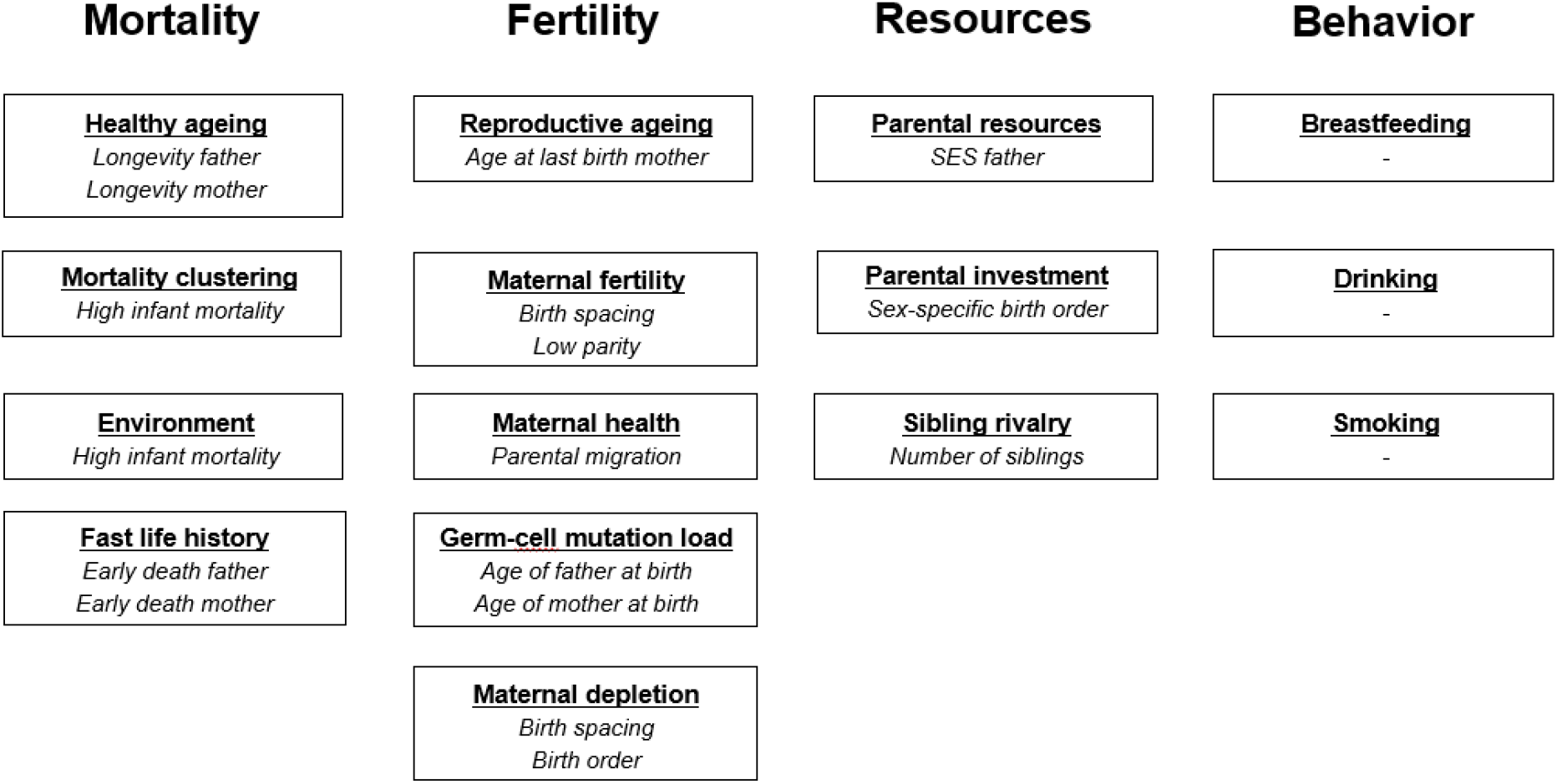
Familial factors associated with offspring survival and their demographic indicators.

## 3. Data & methods

We use LINKS-Zeeland (Mandemakers & Laan, 2017) to study how familial resources affect the relation between parental longevity and offspring survival, defined as age at death or last observation. LINKS-Zeeland is a historical dataset that contains family reconstitutions based on birth, marriage, and death certificates from Zeeland – a coastal province, situated in the southwest of the Netherlands – between 1812 and 1912/1937/1962 respectively (Van den Berg, Van Dijk, Mourits et al., 2018). The size of the dataset makes it possible to make robust estimates of the association between parental longevity and offspring survival, whereas the unique scope of the dataset allows us to test whether the association between parental longevity and offspring survival is dependent on other demographic indicators.

### 3.1 Data selection

To model the association between parental longevity and offspring survival, we selected parent couples for whom the age at death was known. Parental longevity was measured in four different groups: 1. no long-lived parents, 2. a long-lived father, but no long-lived mother, 3. a long-lived mother, but no long-lived father, and 4. two long-lived parents. To be considered long-lived, parents had to belong to the oldest men or women from their birth cohort. We defined longevity as the top 5%, 10%, and 15% survivors in line with earlier research (Van den Berg et al., 2019). We report the top 10% survivors, while the top 5% and top 15% survivors will be presented in supplementary tables. We selected the top percentages of survivors based on Swedish cohort life tables from the Human Mortality Database, as the Swedish life tables are available for the early 19^th^ century and are consistent with lifetables of multiple other industrializing societies at the end of the 19^th^ century (Lindahl-Jacobsen et al., 2013). This procedure has three advantages over making the same selection based on the LINKS dataset itself. First, it produces more accurate estimates of the top survivors. Zeeland was characterized by considerable outmigration and a sizable share of the individuals who survived infancy migrated out of the province. As a result, the top survivors in our dataset are younger than the actual top survivors. Second, the Human Mortality Database has complete cohort life tables for all the parents in our sample, whereas mortality information in LINKS-Zeeland is only available after 1812. Therefore, we do not have to make assumptions on mortality before our observation period. Third, in contrast to LINKS-Zeeland, the Swedish life tables contain enough cases to reliably estimate the upper percentiles of survivors by sex and birth year. Hence, we can more precisely identify the long-lived individuals in our sample.

To test whether the association between parental longevity and offspring survival is affected by other familial factors, we measured high infant mortality in the sibship, familial fertility histories, and shared socioeconomic resources in line with the literature. Infant mortality in the family was measured as the number of offspring dying between the 2^nd^ and 12^th^ month of life. A late-reproducing mother was operationalized as a mother who reproduced after age 45. Low parity families are families with 1, 2, or 3 siblings. Birth intervals are divided in three categories: under 1.5 years, 1.5 to 2.5 years, and more than 2.5 years. Parental migration indicates whether parents migrated within Zeeland after the birth of their first child. Birth order distinguishes between first-, middle-, lastborn children and the rest of the sibship. Paternal socioeconomic status (SES) is measured as the highest social position split into four categories. The elite were fathers who performed learned professions, such as artists, clergymen, doctors, engineers, lawyers, pharmacists, teachers, and veterinarians. Farmers comprise farmers and farm laborers. Middle strata encompass proprietors, managers, clerks, salesmen, and craftsmen. Laborers are all those who performed semi- or unskilled labor. Fathers without a known profession are included as a separate category. Sibling rivalry is operationalized as the number of siblings alive at birth and the number of siblings alive at age 5. Data on parental behavior is not included, as the data LINKS-Zeeland does not contain information on drinking or breastfeeding.

To prevent that certain families were overrepresented in our data, disjoint family trees were selected (see Figure 2). Offspring was included in our sample if their parents married between 1812 and 1862 and the age at death of both parents was known. To identify disjoint families, related family members were removed from this selected sample in two steps. First, we made sure that offspring – the daughters and sons in our sample – did not return as parents. Second, we excluded half-sibs from the data by randomly selecting one childbearing family if fathers or mothers married and reproduced with more than one partner. As shown in Table 1, this left us with data on 16,905 parent couples who had 101,577 children. Only cases with complete information on parental mortality were included in our sample, which also resulted in full information on high infant mortality in the sibship, familial fertility histories, and shared socioeconomic resources. Information on offspring lifespan is available for 81,514 children (80.2%), while information is censored for the other 20,063 children (19.8%). About half of the censored observations are missing from birth, whereas the other half is censored after a vital event, i.e. marriage or childbirth.

**Table 1:** Characteristics of parents and offspring in the studied sample of LINKS Zeeland

|  | Father | Mothers | Sons | Daughters |
| --- | --- | --- | --- | --- |
| **Sample** |  |  |  |  |
| N | 16,905 | 16,905 | 52,367 | 49,210 |
| Birth cohorts | 1741-1842 | 1768-1844 | 1812-1886 | 1812-1885 |
| Age at death |  |  |  |  |
| -available | 16,905 | 16,905 | 41,748 (79.7%) | 39,766 (80.8%) |
| -censored | - | - | 4,318 ( 8.2%) | 4,821 ( 9.8%) |
| -missing | - | - | 6,301 (12.0%) | 4,623 ( 9.4%) |
| **Demographic indicators** |  |  |  |  |
| Age at death* | 62.6 (15.6) | 62.9 (17.4) | 30.0 (33.1) | 32.2 (33.3) |
| Mode lifespan | 71 | 72 | 0 | 0 |
| Age at last birth* | 41.1 (8.1) | 37.5 (5.9) | 39.8 (7.5) | 36.8 (6.2) |
| Number of offspring* | 6.6 (4.0) | 6.3 (3.8) | 6.5 (4.0) | 6.2 (3.8) |
| Birth spacing* | 2.1 (1.3) | 2.1 (1.3) | 2.2 (1.3) | 2.2 (1.3) |
| Migrated within Zeeland | 15.9% | 15.9% | 20.1% | 20.0% |
| Father died before ego was 5* | - | - | 6.3% | 6.0% |
| Mother died before ego was 5* | - | - | 6.2% | 6.2% |
| Age father at birth* | - | - | 34.8 (7.5) | 34.8 (7.5) |
| Age mother at birth* | - | - | 31.8 (6.0) | 31.8 (6.0) |
| Number sibs alive at birth ego* | - | - | 2.4 (2.2) | 2.4 (2.2) |
| Number sibs alive if ego is age 5* | - | - | 3.4 (2.1) | 3.4 (2.2) |

\* Mean + standard deviation between brackets

**Table 2:**
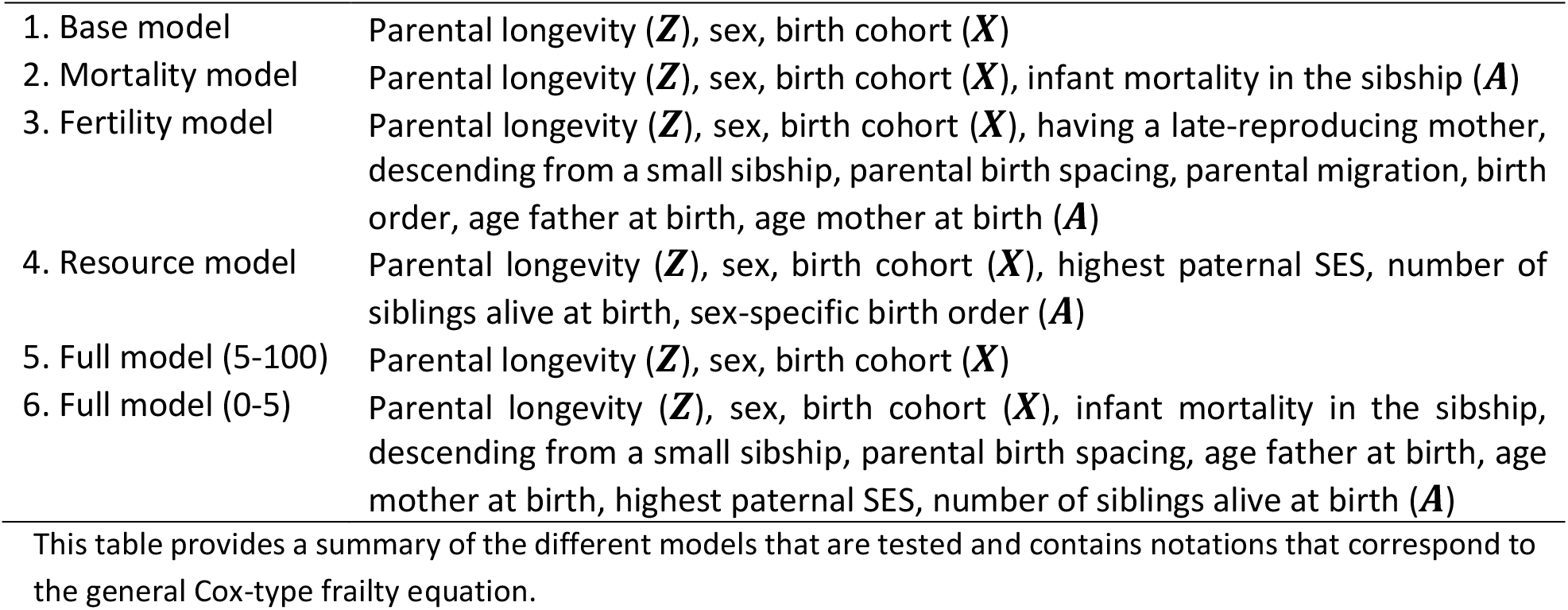
Overview of estimated Cox models

|  |  |
| --- | --- |
| 1. Base model | Parental longevity (**Z**), sex, birth cohort (**X**) |
| 2. Mortality model | Parental longevity (**Z**), sex, birth cohort (**X**), infant mortality in the sibship (**A**) |
| 3. Fertility model | Parental longevity (**Z**), sex, birth cohort (**X**), having a late-reproducing mother, descending from a small sibship, parental birth spacing, parental migration, birth order, age father at birth, age mother at birth (**A**) |
| 4. Resource model | Parental longevity (**Z**), sex, birth cohort (**X**), highest paternal SES, number of siblings alive at birth, sex-specific birth order (**A**) |
| 5. Full model (5-100) | Parental longevity (**Z**), sex, birth cohort (**X**) |
| 6. Full model (0-5) | Parental longevity (**Z**), sex, birth cohort (**X**), infant mortality in the sibship, descending from a small sibship, parental birth spacing, age father at birth, age mother at birth, highest paternal SES, number of siblings alive at birth (**A**) |

This table provides a summary of the different models that are tested and contains notations that correspond to the general Cox-type frailty equation.

**Figure 2:**
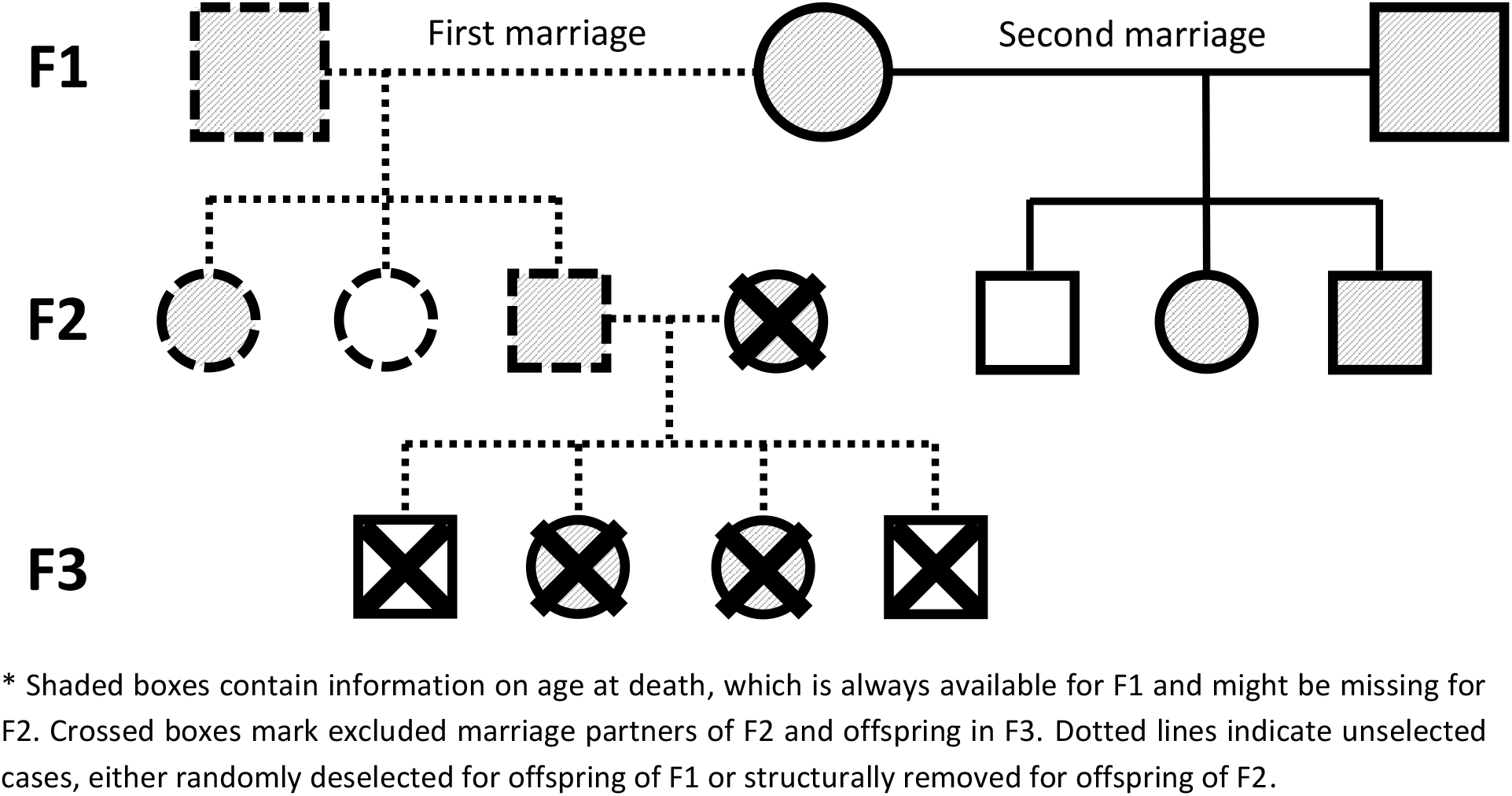
Visualization of disjoint family selection

### 3.2 Sample description

Table 1 shows descriptive statistics on mortality and fertility in Zeeland. The island archipelago was characterized by high infant mortality. 37.6% of all sons and 34.9% of all daughters in our sample died in infancy, which was high in comparison to inland regions in the Netherlands (Hoogerhuis, 2003; Klüsener et al., 2014; Van Dijk & Mandemakers, 2018). The mean lifespan for sons and daughters is also relatively low with 30.0 and 32.2 years, but this is most likely an underestimation due to outmigration (Puschmann, Donrovich & Matthijs, 2017; Van den Berg, Van Dijk, Mourits, et al., 2018), which generally occurred between ages 15 and 50 (Kok, 1997). The mean number of children of 6.5 / 6.3 is comparable to the average fertility in France, Germany, or the Netherlands (Eijkemans et al., 2014), whereas the age at last birth of 36.8 for women and average birth interval of 2.1 are relatively low (Dribe et al., 2017; Eijkemans et al., 2014). The economy was geared towards commercial agriculture. Zeeland specialized in the production of cash crops and grain (Priester, 1998). About two-fifths of the male population worked directly in agriculture either as an agricultural laborer or farmer, while even more unskilled laborers, freighters, and traders were indirectly involved in agriculture (Van Leeuwen & Maas, 2007).

### 3.3 Statistics

The survival of the offspring in our data was studied using Cox regressions with random effects, following previous studies (Dutta et al., 2013; Gudmundsson et al., 2000; Houde et al., 2008; Smith et al., 2009b; Van den Berg et al., 2018; Westendorp et al., 2009). Analyses were done using R version 3.3.0 using the coxme package (R Core Team, 2016; Therneau, 2015). To deal with robustness issues, we censored our analyses at age 100. Earlier studies showed that survival advantages for offspring of long-lived parents can be considered proportional over time (Perls et al., 2002; Van den Berg et al., 2018; Westendorp et al., 2009, Willcox et al., 2006). But since mortality in the first 5 years of life was exceptionally high in Zeeland and up to 40% of all newborns died, we divide our analyses into two parts. The first series of models focuses on offspring survival during childhood, characterized by rapidly receding mortality. We study mortality in the first five years of life with survival censored at age 5 for those living longer. In the second series of models, we focus on the remaining part of the lifespan, during which the individual chances of dying increased exponentially as individuals grow older. In these models, offspring is observed until the end of the observation window or their last observed vital event in Zeeland, usually their death.

In each of the two parts, we estimate five different Cox models to explore whether controlling for other familial resources explains part of the association between parental longevity and offspring survival, defined as age at death or age at last observation in case of censored data. The models are defined as:

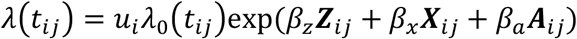

*t*_*ij*_ is the age at death or the age at last observation for child *j* in family *i*. *λ*_0_(*t*_*ij*_) refers to the baseline hazard, which is left unspecified. *u* > 0 refers to an unobserved random effect (frailty) shared by children of a given family. This unobserved heterogeneity shared within sibships was assumed to follow a log-normal distribution. *β*_*z*_ is a vector of regression coefficients for the main effect (***Z***) which corresponds to parental longevity (having a top 10% surviving father, a top 10% surviving mother, or two top 10% surviving parents compared to no parents belonging to the top 10% survivors. *β*_*x*_ contains the regression effects of covariates birth cohort and sex (***X***). *β*_*α*_ contains the regression estimates for a set of extra variables of potential interest (***A***). We estimate four models (model 1-4) for each age group (0-5 years and 5-100 years) and a final model (model 5 and 6) for each age group. The main effects (***Z***) and covariates (***X***) are present in every model, whereas the set of variables (***A***) can differ per model.

To establish whether there is an association between parental longevity and offspring survival, we first estimated the association between parental longevity and offspring survival (*β*_*z*_***Z***) while adjusting for the offspring’s sex and year of birth (*β*_*z*_***X***) in a baseline model (model 1). To test whether this association could be explained by other familial factors, we estimated three different models by adding information on either infant mortality in the sibship (model 2), familial fertility conditions (model 3), or shared socioeconomic resources (model 4) to the baseline model (corresponding to the different variables in ***A***). Model 2 on mortality contains information on parental longevity, offspring’s sex, birth cohort, and infant mortality in the sibship, model 3 on fertility contains information on parental longevity, offspring’s sex, birth cohort, having a late-reproducing mother, descending from a small sibship, parental birth spacing, parental migration, birth order, the age of the father at birth, and age of the mother at birth, while model 4 on socioeconomic models contains information on parental longevity, offspring’s sex, birth cohort, the highest paternal SES, number of siblings alive at birth, and the sex-specific birth order. Finally, variables that associated significantly with offspring survival in any of the previous models were added to a full model (model 5 and 6) that indicates which familial factors had the strongest association with offspring survival between ages 5-100 and 0-5. Variables from the infant mortality, fertility history, and socioeconomic resource models were added to the full model if they were significant with an alpha of 0.0036. This alpha level results from dividing the usual alpha=0.05 by the total number of 14 variables subject to selection. With this approach, when constructing the full models we exclude familial factors that only have a marginal effect on offspring survival but reach statistical significance due to the large sample size of this study.

In our tables, effect sizes will be reported as Exp(β), i.e. Hazard Ratio’s (HR). For model 1-4, confidence intervals and p-values are corrected for the size of the dataset. In the text, we discuss the outcomes of Cox models in terms of HRs and report their 95% confidence intervals in terms of survival advantages. A HR of 0.83, for example, indicates a 17% lower chance of dying and from here we will refer to this as a survival advantage of 17%. We only show abbreviated tables. The full tables are shown in the supplementary Tables A1 and A2 in the appendix. The appendix also includes robustness checks for parental longevity defined as the top 5% and 15% surviving parents in Tables A3 and A4.

## 4. Results

The survival advantages enjoyed by offspring of long-lived parents between ages 5-100 are discussed in paragraph 4.1, while paragraph 4.2 discusses the survival advantages between ages 0-5. In each section, we discuss the results in three steps. First, associations between parental longevity and offspring survival are estimated in a Cox model and controlled for effects of sex and birth cohort. We show cumulative hazard and survival plots for offspring with no parents belonging to the top 10% survivors, a top 10% surviving father, a top 10% surviving mother, and two top 10% surviving parents. Second, we enquire whether the survival advantage enjoyed by offspring of long-lived parents is dependent on other familial resources. We report the HRs for offspring with a top 10% surviving father, a top 10% surviving mother, and two top 10% surviving parents in comparison to offspring with no parents belonging to the top 10% survivors (model 1) while adjusting for effects of high infant mortality in the sibship (model 2), familial fertility histories (model 3), shared socioeconomic resources (model 4), and the three categories combined (model 5). Third, we determine how strong the effect of parental longevity on offspring survival is in comparison to other familial factors between ages 5-100. We show which familial resources are significantly associated with offspring survival and determine how strongly they affect offspring survival in comparison with the effect sizes of having a top 10% surviving father, a top 10% surviving mother, or two top 10% surviving parents.

### 4.1.1 Parental longevity and offspring survival between ages 5-100

We first test whether there is a positive association between parental longevity and offspring survival. Figure 3 shows the cumulative hazard and survival probability between ages 5-100 in panels A and B. Both panels indicate that between ages 5-100, offspring of the top 10% longest-lived fathers and top 10% longest-lived mothers have a similar survival advantage of respectively 17% (HR: 0.83, CI: 0.80-0.87) and 20% (HR: 0.80, CI: 0.77-0.84), over offspring without a long-lived parent. Offspring of two parents belonging to the top 10% survivors enjoyed an even larger survival advantage of 25% (HR: 0.75, CI: 0.69-0.82) over offspring without a long-lived parent. This indicates that the survival advantage of offspring after age 5 is comparable for long-lived mothers and fathers and that the survival advantage increases with the number of long-lived parents.

**Figure 3:**
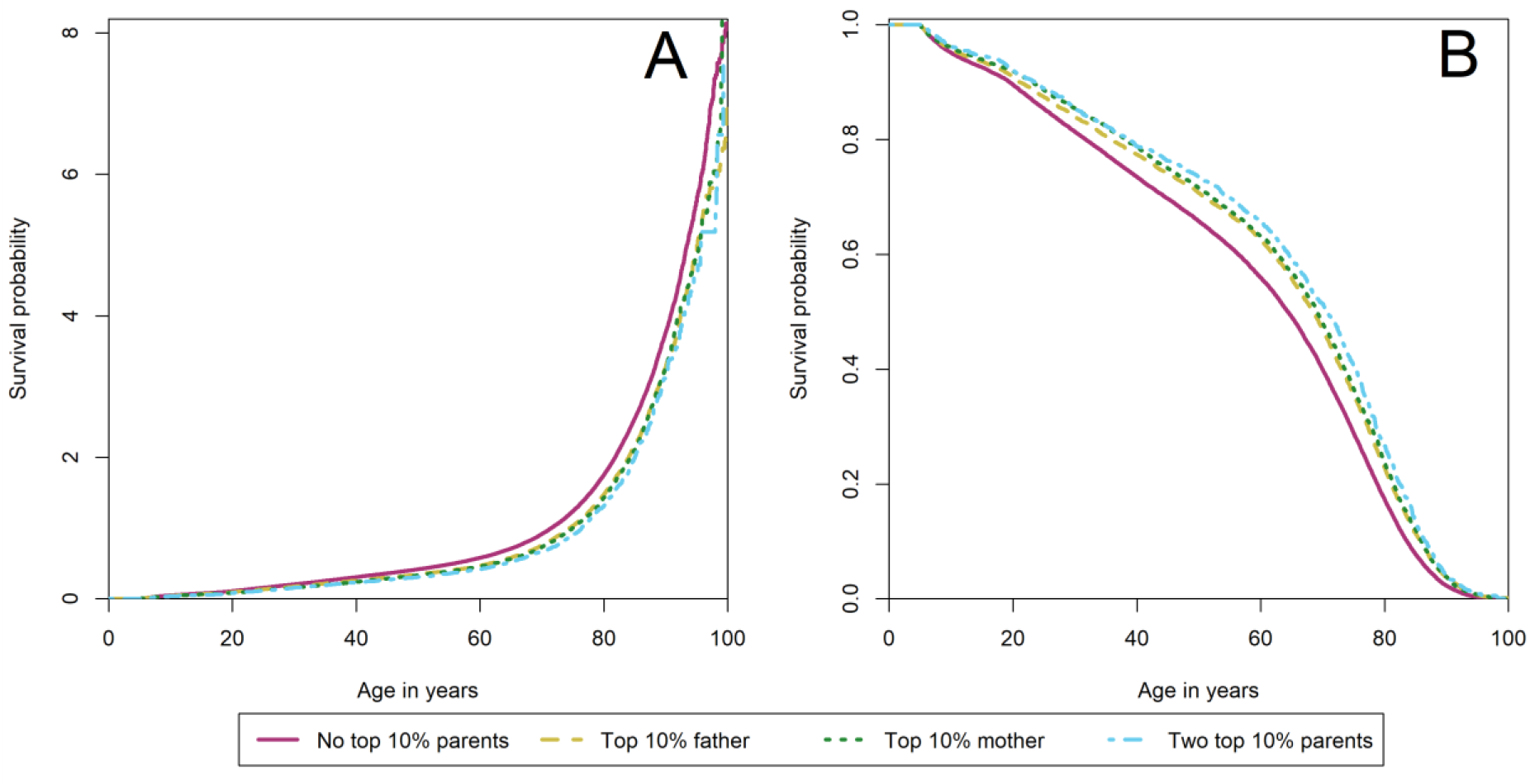
Cumulative hazard (A) and survival plot (B) of the association between having a top 10% surviving parent and offspring survival between ages 5-100. Observations are right-censored for offspring who live past age 100. Cumulative hazard (A) and survival (B) is shown by parental longevity: no top 10% parents (*red, solid line*), having a top 10% father, but no top 10% mother (*yellow, dashed line*), having a top 10% mother, but no top 10% father (*dark green, dotted line*), and having two top 10% parents (*light blue, dotdashed line*). Difference in survival/mortality between the groups represented by the separate lines are formally tested using a cox-type regression analysis. The results are presented in Table 3.

### 4.1.2 Familial factors do not explain the association between parental longevity and offspring survival

Second, we enquire whether the association between parental longevity and increased offspring survival is influenced by effects of high infant mortality in the sibship, familial fertility histories, and shared socioeconomic resources. Table 3 shows the number of observations, number of families, and HR + corrected 95% confidence intervals (CI) for Cox models after controlling for sex and birth year, high infant mortality in the sibship, familial fertility histories, shared socioeconomic resources, and a full model, respectively. Controlling for high infant mortality in the sibship, familial fertility histories and socioeconomic resources did not affect the association between parental longevity and offspring survival between ages 5-100. Survival advantages remained 17% (HR: 0.83, CI: 0.79-0.87) for offspring of top 10% surviving fathers, 20% (HR: 0.80, CI: 0.76-0.85) for offspring of top 10% surviving mothers, and 24% (HR: 0.76, CI: 0.68-0.84) for offspring of two long-lived parents in comparison to offspring without a long-lived parent. Hence, the survival advantage enjoyed by offspring, who died between ages 5-100, of top 10% surviving parents is not explained by infant mortality in the sibship, familial fertility histories of spacing and early reproduction, or better access to socioeconomic resources.

**Table 3:**
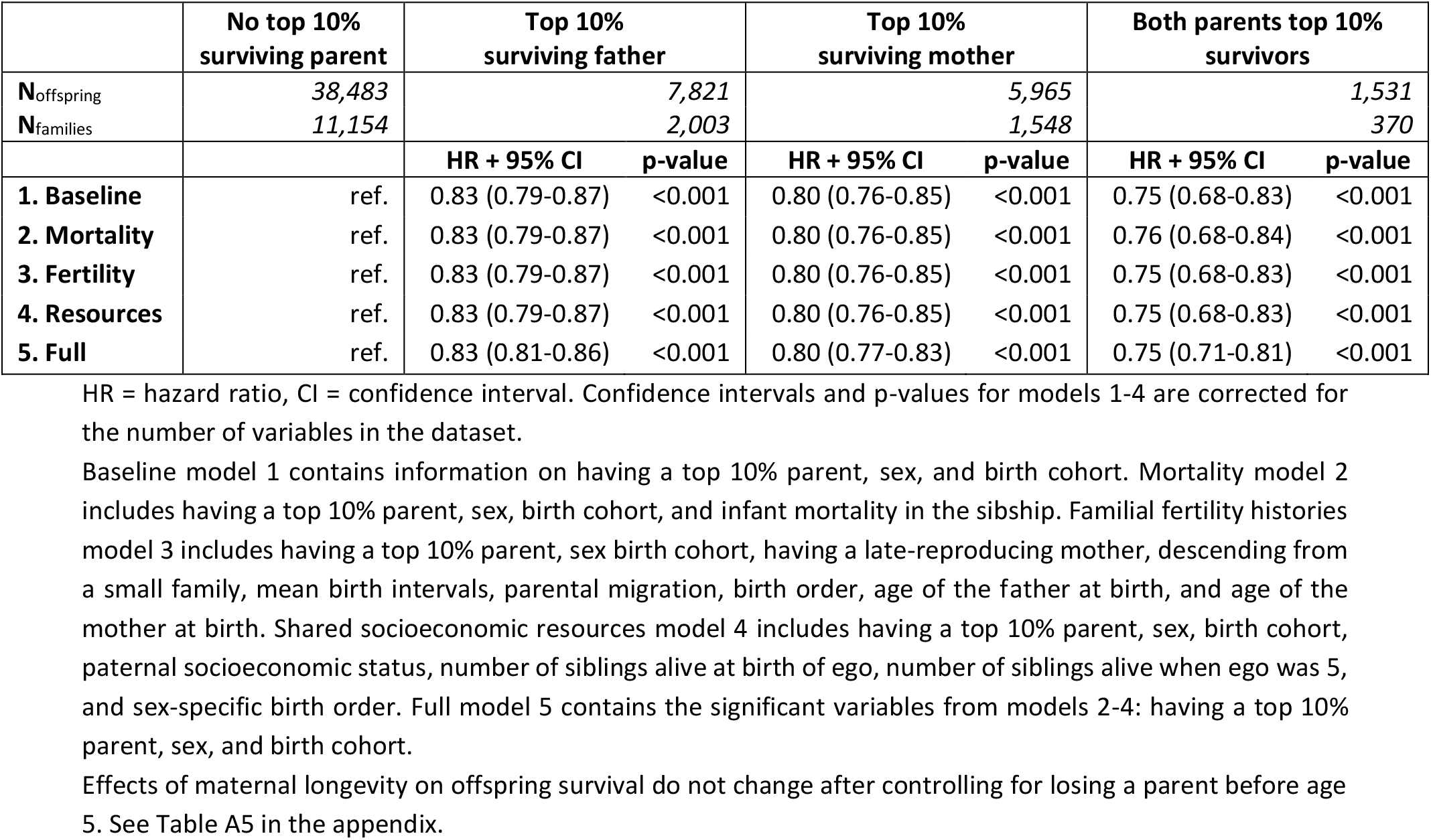
Association between parental longevity and offspring survival between ages 5-100

Robustness checks show that effects of parental longevity on offspring survival do not change after controlling for losing a parent before age 5. See Table A5 in the appendix.

### 4.1.3 Parental longevity is the only important familial factor for survival between ages 5-100

Third, we investigate in the full model (model 5) how the effects of parental longevity compare with other familial factors. This indicates how important parental longevity was between ages 5-100. Table 4 shows the number of observations, number of families, HRs + corrected 95% CI, and corrected p-values.

**Table 4:** Full model (5) of the significant associations between familial resources and offspring survival between ages 5-100

|  | N<sub>offspring</sub> | N<sub>families</sub> | HR + 95% CI | p-value |
| --- | --- | --- | --- | --- |
| **Parental longevity** |  |  |  |  |
| ▪ No top 10% parent | 45,930 | 14,766 | ref. | ref. |
| ▪ Father top 10%, mother not top 10% | 4,294 | 1,072 | 0.83 (0.81-0.86) | <0.001 |
| ▪ Mother top 10%, father not top 10% | 3,236 | 822 | 0.80 (0.77-0.83) | <0.001 |
| ▪ Both parents top 10% | 340 | 79 | 0.75 (0.71-0.81) | <0.001 |

HR = hazard ratio, CI = confidence interval. Reported HRs result from the full model (5).
None of the covariates associated significantly with offspring survival between ages 5-100. Effects of maternal longevity on offspring survival do not change after controlling for losing a parent before age 5. See Table A5 in the appendix.

The full model shows that besides having a top 10% surviving parent, no other variable associated significantly with offspring survival. Hence, survival only increases with each additional parent surviving to the to 10% of their birth cohort.

### 4.2.1 Parental longevity and offspring survival between ages 0-5

Here we focus on survival between ages 0-5. We first test whether there is a positive association between parental longevity and offspring survival. Figure 4 shows that in the first 5 years of life, offspring of fathers belonging to the 10% of their birth cohort had a survival advantage of 8% (HR: 0.92, CI: 0.88-0.97) over offspring without a top 10% surviving parent. Offspring with a top 10% surviving mother, on the other hand, enjoyed a survival advantage of 18% (HR: 0.82, CI: 0.78-0.87) over offspring with no long-lived parents. Offspring of two top 10% surviving parents had a survival advantage of 27% (HR: 0.73, CI: 0.65-0.81) over offspring without a long-lived parent. Thus, having a top 10% surviving mother increases offspring survival with about 20% at every point in the life course, while having a top 10% surviving father will give the same survival benefit though only after 5 years of age. Moreover, similar to the effects after age 5, we observed an increase in offspring survival advantage with the number of long-lived parents.

**Figure 4:**
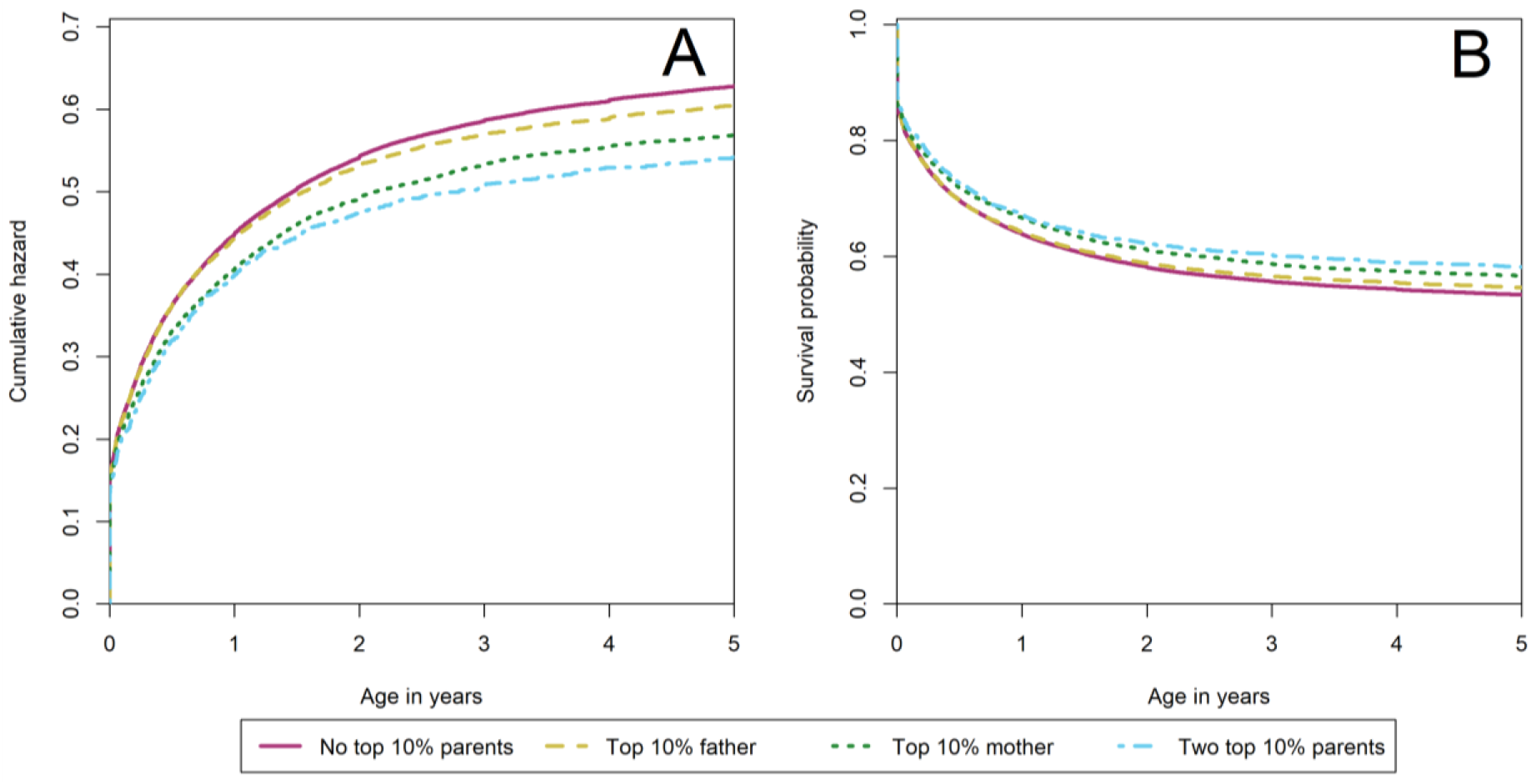
Cumulative hazard (A) and survival plot (B) of the association between having a top 10% surviving parent and offspring survival between ages 0-5. Observations are right-censored for offspring who live past age 5. Cumulative hazard (A) and survival (B) is shown by parental longevity: no top 10% parents (*red, solid line*), having a top 10% father, but no top 10% mother (*yellow, dashed line*), having a top 10% mother, but no top 10% father (*dark green, dotted line*), and having two top 10% parents (*light blue, dotdashed line*). The difference in survival/mortality between the groups represented by the separate lines are formally tested using a cox-type regression analysis. The results are presented in table 5.

### 4.2.2 Familial factors do not explain the association between parental longevity and offspring survival

Second, we enquire whether the association between parental longevity and increased offspring survival is influenced by effects of high infant mortality in the sibship, familial fertility histories, and shared socioeconomic resources. Table 5 shows the association between parental longevity and offspring survival between ages 0-5. Infant mortality in the sibship, familial fertility histories and socioeconomic resources have a marginal impact on the association between parental longevity and offspring survival. The estimated survival advantage of having a long-lived father over having no long-lived parent remained 8% after controlling for high infant mortality, familial fertility histories, or shared socioeconomic resources, but decreased to 7% (HR: 0.93, CI: 0.90-0.96) in the full model. Survival advantages of having a long-lived mother move from 18% (HR: 0.82, CI: 0.77-0.88) to 15% (HR: 0.85, CI: 0.81-0.90) after controlling for infant mortality in the sibship, to 17% (HR: 0.83, CI: 0.78-0.88) after controlling for fertility histories, increases to 19% (HR: 0.81, CI: 0.76-0.87) after controlling for shared socioeconomic resources, and decreases to 15% (HR: 0.85, CI: 0.82-0.89) in the full model. Survival benefits of having two long-lived parents over having no long-lived parent shift from 27% (HR: 0.73, CI: 0.64-0.82) to 24% (HR: 0.76, CI: 0.68-0.85), 26% (HR: 0.74, CI: 0.66-0.84), 28% (HR: 0.72, CI: 0.63-0.81), and 22% (HR: 0.78, CI: 0.72-0.84), respectively. Hence, the association between having a top 10% surviving parent – especially a top 10% surviving mother – and offspring survival advantage between ages 0-5 was also not explained by lower infant mortality in the sibship, familial fertility histories, or shared socioeconomic resources.

**Table 5:** Association between parental longevity and offspring survival between ages 0-5

|  | No top 10%<br>surviving parent | Top 10%<br>surviving father |  | Top 10%<br>surviving mother |  | Both parents top 10%<br>survivors |  |
| --- | --- | --- | --- | --- | --- | --- | --- |
| **N**offspring | 74,336 | 14,381 |  | 10,383 |  | 2,477 |  |
| **N**families | 12,639 | 2,207 |  | 1,665 |  | 394 |  |
|  |  | HR + 95% CI | p-value | HR + 95% CI | p-value | HR + 95% CI | p-value |
| **1. Baseline** | ref. | 0.92 (0.88-0.97) | <0.001 | 0.82 (0.77-0.88) | <0.001 | 0.73 (0.64-0.82) | <0.001 |
| **2. Mortality** | ref. | 0.92 (0.88-0.96) | <0.001 | 0.85 (0.81-0.90) | <0.001 | 0.76 (0.68-0.85) | <0.001 |
| **3. Fertility** | ref. | 0.92 (0.87-0.97) | <0.001 | 0.83 (0.78-0.88) | <0.001 | 0.74 (0.66-0.84) | <0.001 |
| **4. Resources** | ref. | 0.92 (0.87-0.97) | <0.001 | 0.81 (0.76-0.87) | <0.001 | 0.72 (0.63-0.81) | <0.001 |
| **6. Full** | ref. | 0.93 (0.90-0.96) | <0.001 | 0.85 (0.82-0.89) | <0.001 | 0.78 (0.72-0.84) | <0.001 |

HR = hazard ratio, CI = confidence interval. Confidence intervals and p-values for models 1-4 are corrected for the number of variables in the dataset.
Baseline model 1 contains information on having a top 10% parent, sex, and birth cohort. Mortality model 2 includes having a top 10% parent, sex, birth cohort, and infant mortality in the sibship. Familial fertility histories model 3 includes having a top 10% parent, sex birth cohort, having a late-reproducing mother, descending from a small family, mean birth intervals, parental migration, birth order, age of the father at birth, and age of the mother at birth. Shared socioeconomic resources model 4 includes having a top 10% parent, sex, birth cohort, paternal socioeconomic status, number of siblings alive at birth of ego, number of siblings alive when ego was 5, and sex-specific birth order. Full model 6 contains the significant variables from models 2-4: having a top 10% parent, sex, birth cohort, sibling infant mortality, descending from a small sibship, parental birth spacing, age father at birth, age mother at birth, paternal socioeconomic status, and number of siblings alive at birth.

**Table 6:** Full model (6) of the significant associations between familial resources and offspring survival between ages 0-5

|  | N<sub>offspring</sub> | N<sub>families</sub> | HR + 95% CI | p-value |
| --- | --- | --- | --- | --- |
| **Parental longevity** |  |  |  |  |
| ▪ No top 10% parent | 74,336 | 12,639 | ref. | ref. |
| ▪ Father top 10%, mother not top 10% | 14,381 | 2,207 | 0.93 (0.90-0.96) | <0.001 |
| ▪ Mother top 10%, father not top 10% | 10,383 | 1,665 | 0.85 (0.82-0.89) | <0.001 |
| ▪ Both parents top 10% | 2,477 | 394 | 0.78 (0.72-0.84) | <0.001 |
| **Infant mortality** |  |  |  |  |
| Number of infant deaths in the sibship |  |  |  |  |
| ▪ 0 | 34,355 | 8,086 | ref. | ref. |
| ▪ 1 | 28,223 | 4,651 | 1.51 (1.46-1.55) | <0.001 |
| ▪ 2 | 17,273 | 2,176 | 1.82 (1.76-1.88) | <0.001 |
| ▪ 3+ | 21,726 | 2,052 | 2.12 (2.05-2.19) | <0.001 |
| **Familial fertility histories** |  |  |  |  |
| Number of siblings |  |  |  |  |
| ▪ 1 | 1,886 | 1,886 | 1.73 (1.61-1.86) | <0.001 |
| ▪ 2 | 3,392 | 1,696 | 1.54 (1.46-1.63) | <0.001 |
| ▪ 3 | 4,814 | 1,605 | 1.33 (1.26-1.40) | <0.001 |
| ▪ 4+ | 91,485 | 11,718 | ref. | ref. |
| Parental birth spacing |  |  |  |  |
| ▪ <1.5 | 21,972 | 2,978 | ref. | ref. |
| ▪ 1.5-2.5 | 60,948 | 9,979 | 0.81 (0.79-0.84) | <0.001 |
| ▪ >2.5 | 18,657 | 3,948 | 0.71 (0.68-0.74) | <0.001 |
| Age father at birth of ego |  |  |  |  |
| ▪ <25 | 7,698 | - | 0.94 (0.89-0.99) | 0.012 |
| ▪ 25-40 | 70,033 | - | 0.95 (0.92-0.97) | <0.001 |
| ▪ >40 | 23,846 | - | ref. | ref. |
| Age mother at birth of ego |  |  |  |  |
| ▪ <25 | 14,725 | - | 0.86 (0.82-0.90) | <0.001 |
| ▪ 25-40 | 76,526 | - | 0.95 (0.92-0.98) | 0.004 |
| ▪ >40 | 10,326 | - | ref. | ref. |
| **Shared socioeconomic resources** |  |  |  |  |
| Highest socioeconomic status father |  |  |  |  |
| ▪ *Farmers* | 18,162 | 2,597 | 0.74 (0.72-0.77) | <0.001 |
| ▪ *Elite* | 2,013 | 308 | 1.08 (1.00-1.17) | 0.048 |
| ▪ *Middle strata* | 33,872 | 5,498 | 0.99 (0.96-1.01) | 0.252 |
| ▪ *Laborers* | 46,655 | 8,215 | ref. | ref. |
| ▪ *NA* | 875 | 287 | 0.82 (0.72-0.92) | <0.001 |
| Number of living siblings at birth |  |  |  |  |
| ▪ 0-1 | 43,222 | - | 0.85 (0.82-0.88) | <0.001 |
| ▪ 2-4 | 41,534 | - | 0.95 (0.92-0.97) | <0.001 |
| ▪ 5+ | 16,821 | - | ref. | ref. |

HR = hazard ratio, CI = confidence interval. Reported HRs result from the full model (5) and confidence intervals and p-values are corrected for the number of variables in the dataset.
Effects of having a late-reproducing mother, birth order, and sex-specific birth order were not significant.
Short and long birth intervals occurred both in small and large families. Moreover, there was no interaction between sibship size and birth spacing. See Table A7 in the appendix.

Robustness checks show that effects of parental longevity on offspring survival do not disappear after controlling for losing a parent before age 5. See Table A6 in the appendix. However, between ages 0-5 the survival advantages enjoyed by offspring with a long-lived father, mother, or two long-lived parents decrease from 7% (HR: 0.93, CI: 0.90-0.96) to 6% (HR: 0.94, CI: 0.91-0.97), from 15% (HR: 0.85, CI: 0.82-0.89) to 11% (HR: 0.89, CI: 0.85-0.92), and from 22% (HR: 0.78, CI: 0.72-0.84) to 18% (HR: 0.82, CI: 0.76-0.88).

### 4.2.3 Before age 5, infant mortality and fertility histories have a stronger association with offspring survival than maternal longevity

Last, we investigate the full model (model 6) and compare the effect size of parental longevity with the maximum effect size of other familial factors to indicate how important parental longevity was between ages 0-5. It should be noted that survival advantages cannot be directly compared to survival disadvantages, as the scores take place on a different scale. Survival advantages run on a scale from 0% to 100%, whereas survival disadvantages run on a scale from 0% to infinity. However, survival advantages can easily be transformed into a percentage of decreased survival disadvantages, by dividing 1 by the HR. For example, offspring of long-lived fathers have a HR of 0.93, which corresponds with a decreased survival disadvantage of 1 / 0.93 = 1.08, i.e. a decreased survival disadvantage of 8%. Calculating these scores makes it possible to compare survival disadvantages to the survival advantages enjoyed by offspring of long-lived parents. Hence, survival advantages of 7%, 15%, and 22% for offspring of long-lived fathers, mothers, and two long-lived parents, correspond with decreased survival disadvantages of 8%, 18%, and 28%, respectively.

The full models show that before the age of 5, the decreased survival disadvantage of having a top 10% surviving parent was relatively small in comparison to the increased survival disadvantage of high infant mortality in the sibship, familial fertility histories, and shared socioeconomic resources. High infant mortality in the sibship had the largest effect size on offspring survival. Offspring had a survival disadvantage of 51% (HR: 1.51, CI: 1.46-1.55) if one sibling died during infancy. This survival disadvantage was 82% (HR: 1.82, CI: 1.76-1.88) if two siblings died during infancy and 112% (HR: 2.12, CI: 2.05-2.19) if three or more siblings died during infancy. Hence, infant mortality in the sibships and parental longevity are two independent factors, of which high infant mortality in the sibship had a larger effect on survival before age 5 than maternal longevity.

Familial fertility histories had robust associations with offspring survival before age 5. Descending from a small family associated with a survival disadvantage of 73% (HR: 1.73, CI: 1.61-1.86) in single-child families, 54% (HR: 1.54, CI: 1.46-1.63) for families with two children, and 33% (HR: 1.33, CI: 1.26-1.40) for families with three children. Offspring whose parents had long or medium birth intervals had a decreased survival advantage of 41% (HR: 0.71, CI: 0.68-0.74) and 23% (HR: 0.81, CI: 0.79-0.84), respectively, compared to offspring whose parents had short birth intervals. Offspring whose mother was younger than 25 at the time of their birth had a decreased survival disadvantage of 16% (HR: 0.86, CI: 0.82-0.90) compared to offspring whose mother was over 40 years old, while offspring with mothers between 25 and 40 at the time of their own birth had a survival advantage of 5% (HR: 0.95, CI: 0.92-0.98). Offspring with a father between ages 25 and 40 at birth had a decreased survival disadvantage of 5% (HR: 0.95, CI: 0.92-0.97) over offspring with a father who was over 40 years old at birth. Offspring whose father was younger than 25 years had a similar decreased survival disadvantage of 6% (HR: 0.94, CI: 0.89-0.99) over offspring with a father between ages 25 and 40 at birth. Firstborn offspring initially had a survival disadvantage compared to other offspring, but this effect disappeared after we controlled for high infant mortality in the sibship and shared socioeconomic resources (see supplementary table A2). Thus, descending from a small family, parental birth intervals, mother’s age at birth, and father’s age at birth affect survival between ages 0-5 independently from parental longevity. Of these effects, small family size and birth intervals longer than 2.5 years had a larger effect on offspring survival before age 5 than having a top 10% surviving mother.

Shared socioeconomic resources had relatively weak effects on offspring survival. Offspring of farmers had a decreased survival disadvantage of 35% (HR: 0.74, CI: 0.72-0.77) and offspring of the elite had a, barely significant, survival disadvantage of 8% (CI: 1.08, CI: 1.00-1.17). Having fewer living siblings at birth associated with a decreased survival disadvantage of 18% (HR: 0.85, CI: 0.82-0.88) for offspring with 0-1 siblings alive at birth and 5% (HR: 0.95, CI: 0.92-0.97) for offspring with 2-4 siblings at birth in comparison to offspring with 5 or more siblings alive at birth. Hence, offspring of farmers had a stronger survival advantage before age 5 than offspring of top 10% surviving mothers, whereas the maximum effect of sibling rivalry was similar to the effect of having a top 10% surviving father.

## 5. Discussion

In this paper we set out to investigate whether the intergenerational transmission of longevity was affected by other familial factors than familial longevity. We revisit the question asked by Gavrilov & Gavrilova (2015) and You, Gi & Yi (2010) with extensive data on familial mortality that contains longitudinal information on familial resources. Using newly available demographic data, parental longevity, offspring survival, high infant mortality in the sibship, familial fertility history, and shared socioeconomic resources were associated with offspring survival for 16,905 disjoint families. This sample from LINKS (Mandemakers & Laan, 2017) is unique in terms of sample size, available demographic information, and observation period, enabling us to follow offspring survival for 101,577 children from 16,905 disjoint families with information on parental longevity and a wide range of other familial resources. By testing effects of parental longevity with other familial factors, such as infant mortality in the sibship, descending from a small sibship, birth spacing, parental ages at birth, paternal socioeconomic status, and sibling rivalry, we were able to determine whether the beneficial effect of having long-lived parents was dependent on other familial resources.

We improved on the earlier studies in multiple ways. The major strength of our analysis rests in the scope and range of our dataset. The used sample of available families and individuals within these families is much larger than in previous studies (Dutta et al., 2013; Gavrilov & Gavrilova, 2015; Gudmundsson et al., 2000; Houde et al., 2008; Smith et al., 2009b; Van den Berg et al., 2018; Westendorp et al., 2009; You, Gi & Yi, 2010). This allowed us to simultaneously test a wide range of hypotheses and correct for effects of multiple testing. Second, we applied a more robust definition of longevity that is not affected by sex-differences in lifespan or incremental increases in survival over time (Van den Berg et al., 2019). Longevity is defined as a top percentage of the general population, rather than a share of the oldest individuals in our dataset. Third, we used multiple cut-offs to define parental longevity. This allowed us to verify that our results were not dependent on our definition of paternal longevity. Moreover, by keeping the contrasts between groups constant, we showed that longevity was actually present for the entire top 10% and not for a smaller contingent of long-lived parents. Fourth, rather than testing whether long-lived individuals were more likely to have long-lived parents, we focused on the entire lifespan for all offspring of long-lived individuals, because increased survival in the offspring of long-lived individuals is indicative of a transmission of parental longevity and survival advantages for offspring of long-lived persons are life-long sustained (Perls et al., 2002; Van den Berg et al., 2018; Westendorp et al., 2009, Willcox et al., 2006). This allowed us to not only enquire whether certain characteristics are more common in long-lived individuals, but to also test whether sibling characteristics, parental characteristics, and family compositions affected survival into extreme ages for entire sibships. In summary, the focus on the family and observation from cradle to the grave allowed us to have more information on familial resources. Therefore, we were able to show that the association between parental longevity and offspring survival in Zeeland was independent of a wide range of familial factors.

Parental longevity provided a survival benefit of about 20% for offspring between ages 5-100. Between ages 0-5 this effect is similar for offspring of long-lived mothers and somewhat weaker for offspring of long-lived fathers. The association between parental longevity and offspring survival was not affected by other familial factors. We report no significant survival differences between offspring of long-lived mothers and offspring of long-lived fathers between ages 5-100 but we did observe such effects at ages before 5 years. That the mother has the potency to become long-lived might have been especially important in the first years of life, as healthy mothers can provide their offspring with survival advantages in the womb or postnatally, for example by breastfeeding. Giving birth to children with a higher birth weight can make offspring less susceptible to infectious disease and more likely to recover from food or water poisoning. In Zeeland the first 5 years of life were characterized by exceptionally high mortality (Van Poppel, Jonker & Mandemakers, 2005; Klüsener et al., 2014). In an environment where one in three children did not live to be 5 years old, every survival advantage counted. Therefore, the beneficial effect of having a mother with longevity potential compared to the effect of having a father with longevity potential might have been emphasized in our study. Further study is required to understand how high mortality regimes affect the association between parental longevity and offspring survival.

In earlier studies, stronger maternal lifespan and longevity effects on offspring survival were found (Bocquet-Appel & Jakobi, 1990; Kemkes-Grottenthaler, 2004; Kerber et al., 2001; Piraino et al., 2014; Salaris, Tedesco & Poulain, 2013; Van den Berg et al., 2018; Westendorp & Kirkwood, 2001). These studies offered possible explanations of longevity being transmitted by mitochondrial DNA (mtDNA) since offspring obtain mtDNA only through mothers (Van den Berg et al., 2018). Offspring of long-lived mothers in Zeeland had a survival advantage over offspring of long-lived fathers and non-long-lived parents only during the initial five years of life, also in a subsample where offspring did not lose their parent. This may indicate that mitochondrial functions – the energy regulator of the cell – contribute to longevity mainly by early developmental benefit, but also hints at the importance of having a healthy mother for in-utero development in a high mortality environment (Floud et al., 2011; Van den Berg et al., 2018; Wrigley, 2004).

Between ages 5 and 100, parental longevity was by the only predictor of offspring survival. Contrary to earlier findings in the literature, we found no evidence that offspring survival between ages 5-100 was affected by high mortality in the family, family fertility histories, or socioeconomic resources. In the literature, the enduring effects of high sibling infant mortality on individual survival are well-documented for Southern Sweden (Bengtsson & Lindström, 2000, 2003; Quaranta, 2013) and have recently been replicated for the Netherlands and Utah (Van Dijk, Janssens & Smith, 2018). High sibling infant mortality indicates that there might be something going structurally ‘wrong’ in these families, for example genetic defects, extremely unhealthy environments, or behavior (Van den Boomen & Ekamper, 2015; Hedefalk, Quaranta & Bengtsson, 2017; Van Dijk & Mandemakers, 2018; Walhout, 2019). However, in our study the association between infant mortality in the sibship and offspring survival was insignificant after controlling for the size of the dataset, indicating that infant mortality in the sibship had only a minor impact on individual chances to become long-lived. Associations between family fertility histories and offspring survival have been less thoroughly studied. Hin, Ogórek, and Hedefalk (2016) reported that having a late-reproducing mother or fewer siblings associated with increased survival after age 50. However, we found no evidence that having a late-reproducing mother, lower birth order, longer birth intervals, or younger parents associated with offspring survival associated with offspring survival between ages 5-100. Factors associated with family fertility histories seem to affect survival before age 5, but afterwards were marginal at best. Finally, in line with most other studies on mortality in the 19^th^ and early 20^th^ century, we found no socioeconomic gradient in mortality after age 50 (Bengtsson & Van Poppel, 2011; Edvinsson & Broström, 2012). Generally, social gradients in longevity did not appear until after the 1950s, and sometimes even later (Debiasi & Dribe, 2019; Edvinsson & Broström, 2017; Smith et al., 2009b; Temby & Smith, 2014). Accordingly, effects of sibling infant mortality, family fertility histories, and socioeconomic resources on individual chances to become long-lived were marginal at best.

Before age 5, offspring of long-lived fathers had a smaller survival advantage of 7%. The survival advantage enjoyed by offspring of long-lived mothers remained roughly 20%, but was modest in comparison to known additive effects of high infant mortality in the sibship (Broström, Edvinsson & Engberg, 2018; Gagnon et al., 2009; Hin, Ogórek & Hedefalk, 2016; Janssens, Messelink & Need, 2010; Sommerseth, 2018; Van Dijk & Mandemakers, 2018), descending from a small family (Doblhammer & Oeppen, 2003), parental birth spacing (Dewey & Cohen, 2007; Kozuki et al., 2013), or having a farming father (Breschi et al., 2011; Edvinsson et al., 2005; Janssens & Pelzer, 2012; Schumacher & Oris, 2011; Van Poppel, Jonker & Mandemakers, 2005). However, the survival advantage enjoyed by offspring of long-lived parents is indicative of one of the mechanisms behind child mortality. In total, we can distinguish three different mechanisms that affect offspring survival in the first five years of life. First, there might be some inherited frailty, as offspring of parents who died in early adulthood and individuals from small sibling sets have higher mortality rates. Inherited frailty seems to be the strongest predictor of child mortality. Second, there is the importance of parental care and socioeconomic resources: not losing a parent, having a farmer as a father or fewer siblings at birth increase survival. Third, there is the importance of maternal health or maternal genetic influence on development: long-lived mothers, families with longer birth spacing, and younger mothers produce offspring that lives longer. Effects of maternal health are about as strong as the effects of parental care and socioeconomic resources. Hence, we provided additional evidence for the importance of familial longevity on child mortality.

Our findings indicate some fruitful areas for further research. First, our results suggest that the association between parental longevity and increased offspring survival is not affected by other familial factors. However, this is not necessarily the case for other measures of familial clustering, as parental longevity is just one indicator of familial longevity. Second, parental behaviors such as breastfeeding practices, daily diets, and alcohol consumption are known determinants of offspring survival in early life and may also affect survival in later life (Black et al., 2008; Cnattingius et al., 1998; Hill et al., 2000; Huizink & Mulder, 2006; Ji et al., 1997; van den Boomen & Ekamper, 2015; Walhout, 2010). We were not able to control for these factors, as historical databases generally do not contain information on behavior. Studies on contemporary populations are necessary to indicate whether behavior can affect the association between parental longevity and offspring survival. Third, little is known about the effect of long-distance migration has on the intergenerational transmission of longevity. We found no difference in the association between parental longevity and offspring between stayers and migrants within Zeeland. However, this does not mean that individuals who migrated to a radically new environment with a different disease environment, new social customs, and less social support enjoyed the same survival advantages as their siblings. Fourth, it should be noted that we studied a historical population and some of our effects are known to be subject to changes over time. For example, socioeconomic effects on differences in survival have most likely increased over time, as they were weak at best in the 19^th^ century (Bengtsson & van Poppel, 2011; Edvinsson & Broström, 2012; Edvinsson & Lindkvist, 2011). Today, socioeconomic effects on differences in survival are almost axiomatic and more connected to education and lifestyle rather than access to food (Edvinsson & Broström, 2012; Elo, 2009; Mackenbach et al., 2008), indicating that parental socioeconomic status had a different effect on offspring survival in the past than today (Clouston et al., 2016; Debiasi & Dribe, 2019; Edvinsson & Broström, 2012). There are indications that today the association between parental longevity and offspring survival is affected by socioeconomic status (Temby & Smith, 2014). Understanding when and why this interplay between parental social position and parental longevity occurs can give us better insight in the mechanisms behind familial clustering of longevity. Finally, levels of child mortality have decreased dramatically since the 1880s (Human Mortality Database, 2018) and nowadays have almost no impact on an individual’s chances to become long-lived. Yet, the survival advantage that offspring of long-lived mother enjoy in early life is indicative of underlying biological mechanisms that probably still affect survival today, both early and later in life.

Parental longevity was the most important familial resource for offspring survival at every year in life between 1812 and 1962. Its beneficial effect on offspring survival is transmitted equally by both fathers and mothers, although the beneficial effect of having a long-lived father mainly starts after the age of 5 years. This emphasizes the importance of studying the timing of lifespan advantages, especially since today’s (super)centenarians were born when a significant share of the population still died in the first 5 years of life. Using the LINKS data, the effects of social and behavioral factors on offspring longevity were extensively studied. Infant mortality in the sibship, familial fertility histories, and shared socioeconomic resources did not affect the association between parental longevity and offspring survival. Therefore, we conclude that future research should focus on better characterization of the social and behavioral effects of familial longevity, as well as on the mapping of the genetic contribution to familial longevity.

## Acknowledgements

This project was supported by the NOW under grant 360-53-180.

## 7. Appendix

**Table A1:**
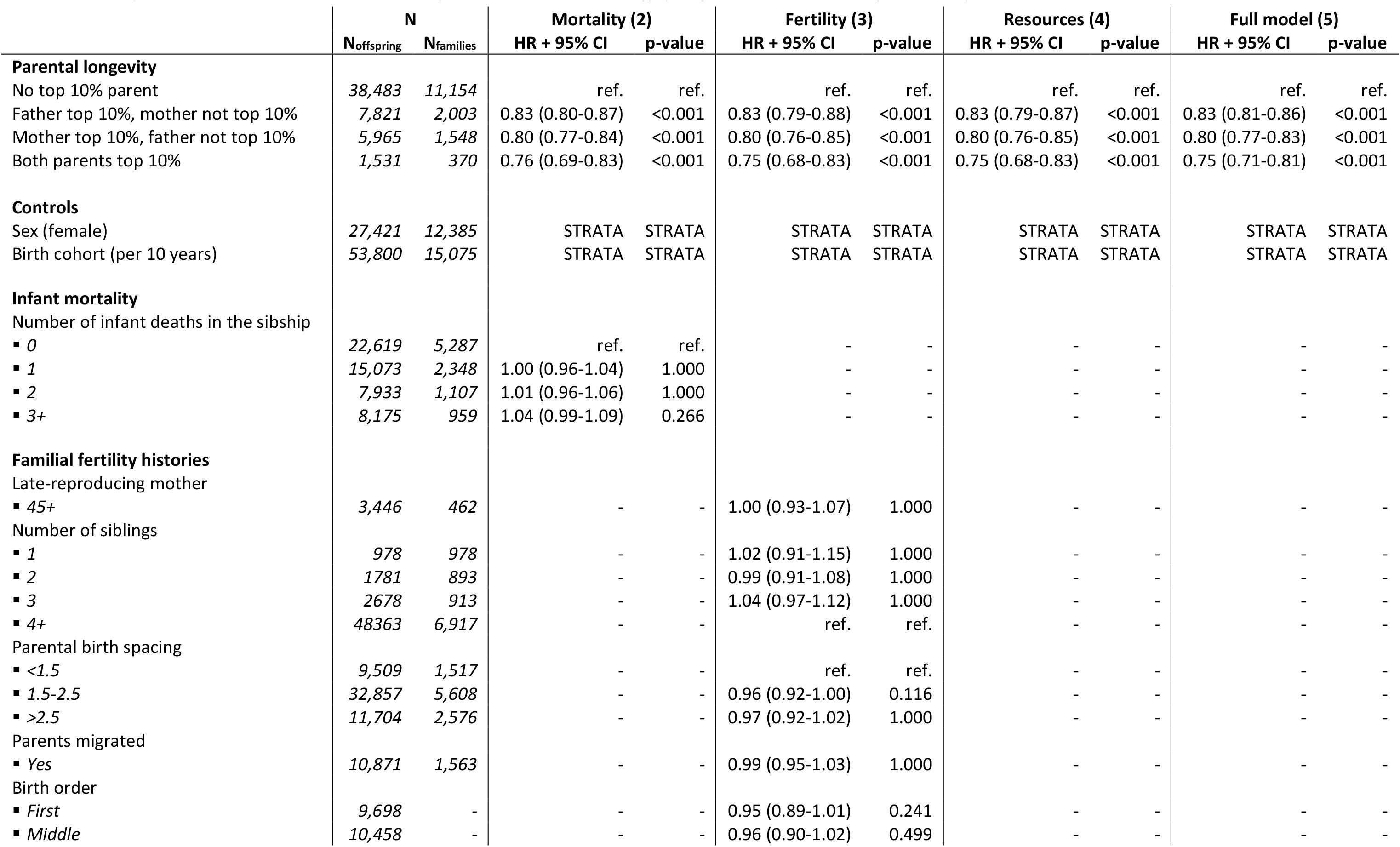
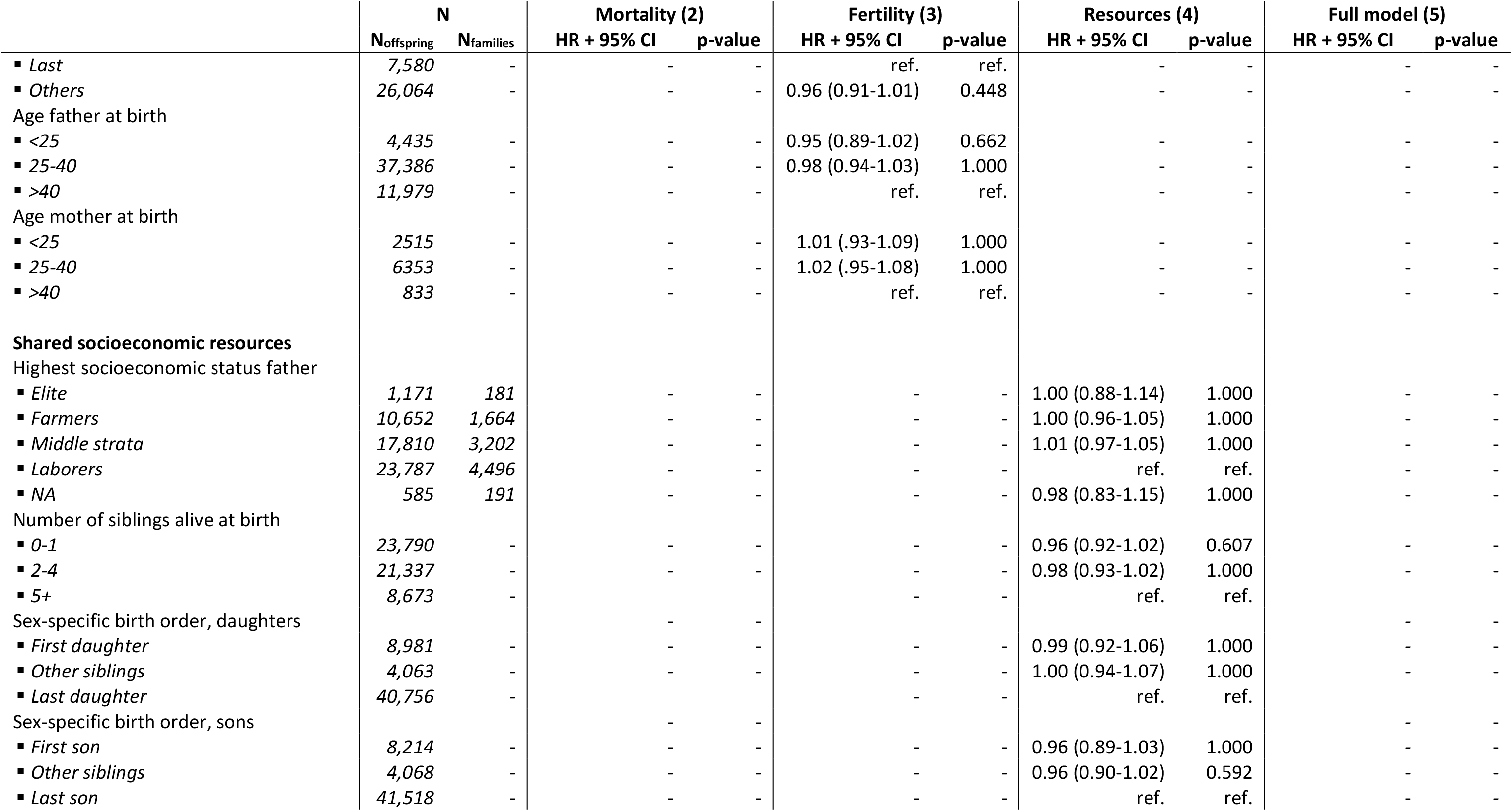
Separate model associations between familial resources and offspring survival between ages 5-100, full table

**Table A2:** Separate model associations between familial resources and offspring survival between ages 0-5, full table

|  | N |  | Mortality model (2) |  | Fertility model (3) |  | Resources model (4) |  | Full model (6) |  |
| --- | --- | --- | --- | --- | --- | --- | --- | --- | --- | --- |
|  | N<sub>offspring</sub> | N<sub>families</sub> | HR + 95% CI | p-value | HR + 95% CI | p-value | HR + 95% CI | p-value | HR + 95% CI | p-value |
| **Parental longevity** |  |  |  |  |  |  |  |  |  |  |
| No top 10% parent | 74,336 | 12,639 | ref. | ref. | ref. | ref. | ref. | ref. | ref. | ref. |
| Father top 10%, mother not top 10% | 14,381 | 2,207 | 0.92 (0.88-0.96) | <0.001 | 0.92 (0.87-0.97) | <0.001 | 0.92 (0.87-0.97) | <0.001 | 0.93 (0.90-0.96) | <0.001 |
| Mother top 10%, father not top 10% | 10,383 | 1,665 | 0.85 (0.81-0.90) | <0.001 | 0.83 (0.78-0.88) | <0.001 | 0.81 (0.76-0.87) | <0.001 | 0.85 (0.82-0.89) | <0.001 |
| Both parents top 10% | 2,477 | 394 | 0.76 (0.68-0.85) | <0.001 | 0.74 (0.66-0.84) | <0.001 | 0.72 (0.63-0.81) | <0.001 | 0.78 (0.72-0.84) | <0.001 |
| **Controls** |  |  |  |  |  |  |  |  |  |  |
| Sex (female) | 49,210 | - | 0.83 (0.80-0.85) | <0.001 | 0.82 (0.80-0.85) | <0.001 | 0.83 (0.80-0.86) | <0.001 | 0.83 (0.81-0.84) | <0.001 |
| Birth cohort (per 10 years) | 101,577 | 16,905 | 1.05 (1.04-1.06) | <0.001 | 1.04 (1.03-1.05) | <0.001 | 1.03 (1.02-1.04) | <0.001 | 1.02 (1.01-1.03) | <0.001 |
| **Infant mortality** |  |  |  |  |  |  |  |  |  |  |
| Number of infant deaths in the sibship |  |  |  |  |  |  |  |  |  |  |
| ▪ 0 | 34,355 | 8,086 | ref. | ref. | - | - | - | - | ref. | ref. |
| ▪ 1 | 28,223 | 4,651 | 1.46 (1.40-1.52) | <0.001 | - | - | - | - | 1.51 (1.46-1.55) | <0.001 |
| ▪ 2 | 17,273 | 2,176 | 1.75 (1.67-1.84) | <0.001 | - | - | - | - | 1.82 (1.76-1.88) | <0.001 |
| ▪ 3+ | 21,726 | 2,052 | 2.09 (2.00-2.19) | <0.001 | - | - | - | - | 2.12 (2.05-2.19) | <0.001 |
| **Familial fertility histories** |  |  |  |  |  |  |  |  |  |  |
| Late-reproducing mother |  |  |  |  |  |  |  |  |  |  |
| ▪ 45+ | 6,215 | 743 | - | - | 0.96 (0.89-1.04) | 1.000 | - | - | - | - |
| Number of siblings |  |  |  |  |  |  |  |  |  |  |
| ▪ 1 | 1,886 | 1,886 | - | - | 1.21 (1.08-1.35) | <0.001 | - | - | 1.73 (1.61-1.86) | <0.001 |
| ▪ 2 | 3,392 | 1,696 | - | - | 1.12 (1.03-1.23) | <0.001 | - | - | 1.54 (1.46-1.63) | <0.001 |
| ▪ 3 | 4,814 | 1,605 | - | - | 1.04 (0.97-1.12) | 1.000 | - | - | 1.33 (1.26-1.40) | <0.001 |
| ▪ 4+ | 91,485 | 11,718 | - | - | ref. | ref. | - | - | ref. | ref. |
| Parental birth spacing |  |  |  |  |  |  |  |  |  |  |
| ▪ <1.5 | 21,972 | 2,978 | - | - | ref. | ref. | - | - | ref. | ref. |
| ▪ 1.5-2.5 | 60,948 | 9,979 | - | - | 0.74 (0.71-0.77) | <0.001 | - | - | 0.81 (0.79-0.84) | <0.001 |
| ▪ >2.5 | 18,657 | 3,948 | - | - | 0.56 (0.53-0.59) | <0.001 | - | - | 0.71 (0.68-0.74) | <0.001 |
| Parents migrated |  |  |  |  |  |  |  |  |  |  |
| ▪ Yes | 20,349 | 2,683 | - | - | 0.97 (0.93-1.02) | 1.000 | - | - | - | - |
| Birth order |  |  |  |  |  |  |  |  |  |  |
| ▪ First | 16,901 | - | - | - | 0.92 (0.87-0.98) | <0.001 | - | - | - | - |
| ▪ Middle | 19,486 | - | - | - | 0.98 (0.93-1.03) | 1.000 | - | - | - | - |
|  | N<sub>offspring</sub> | N<sub>families</sub> | HR + 95% CI | p-value | HR + 95% CI | p-value | HR + 95% CI | p-value | HR + 95% CI | p-value |
| ▪ *Last* | 15,010 | - | - | - | ref. | ref. | - | - | - | - |
| ▪ *Others* | 50,180 | - | - | - | 0.98 (0.93-1.02) | 1.000 | - | - | - | - |
| Age father at birth |  |  |  |  |  |  |  |  |  |  |
| ▪ <25 | 7,698 | - | - | - | 0.92 (0.85-0.99) | 0.007 | - | - | 0.94 (0.89-0.99) | 0.012 |
| ▪ 25-40 | 70,033 | - | - | - | 0.94 (0.90-0.97) | <0.001 | - | - | 0.95 (0.92-0.97) | <0.001 |
| ▪ >40 | 23,846 | - | - | - | ref. | ref. | - | - | ref. | ref. |
| Age mother at birth |  |  |  |  |  |  |  |  |  |  |
| ▪ <25 | 14,725 | - | - | - | 0.83 (0.77-0.89) | <0.001 | - | - | 0.86 (0.82-0.90) | <0.001 |
| ▪ 25-40 | 76,526 | - | - | - | 0.93 (0.88-0.98) | 0.002 | - | - | 0.95 (0.92-0.98) | 0.004 |
| ▪ >40 | 10,326 | - | - | - | ref. | ref. | - | - | ref. | ref. |
| **Shared socioeconomic resources** |  |  |  |  |  |  |  |  |  |  |
| Highest socioeconomic status father |  |  |  |  |  |  |  |  |  |  |
| ▪ *Elite* | 2,013 | 308 | - | - | - | - | 1.09 (0.95-1.24) | 0.906 | 1.08 (1.00-1.17) | 0.048 |
| ▪ *Farmers* | 18,162 | 2,597 | - | - | - | - | 0.78 (0.74-0.82) | <0.001 | 0.74 (0.72-0.77) | <0.001 |
| ▪ *Middle strata* | 33,872 | 5,498 | - | - | - | - | 0.98 (0.94-1.02) | 1.000 | 0.99 (0.96-1.01) | 0.252 |
| ▪ *Laborers* | 46,655 | 8,215 | - | - | - | - | ref. | ref. | ref. | ref. |
| ▪ *NA* | 875 | 287 | - | - | - | - | 0.86 (0.71-1.05) | 0.383 | 0.82 (0.72-0.92) | <0.001 |
| Number of siblings alive at birth |  |  |  |  |  |  |  |  |  |  |
| ▪ 0-1 | 43,222 | - | - | - | - | - | 0.76 (0.72-0.79) | <0.001 | 0.85 (0.82-0.88) | <0.001 |
| ▪ 2-4 | 41,534 | - | - | - | - | - | 0.86 (0.83-0.90) | <0.001 | 0.95 (0.92-0.97) | <0.001 |
| ▪ 5+ | 16,821 | - | - | - | - | - | ref. | ref. | ref. | ref. |
| Sex-specific birth order, daughters |  |  |  |  |  |  |  |  |  |  |
| ▪ *First daughter* | 14,975 | - | - | - | - | - | 0.95 (0.89-1.02) | 0.404 | - | - |
| ▪ *Other siblings* | 79,054 | - | - | - | - | - | 1.00 (0.94-1.05) | 1.000 | - | - |
| ▪ *Last daughter* | 7,548 | - | - | - | - | - | ref. | ref. | - | - |
| Sex-specific birth order, sons |  |  |  |  |  |  |  |  |  |  |
| ▪ *First son* | 15,258 | - | - | - | - | - | 0.98 (0.92-1.05) | 1.000 | - | - |
| ▪ *Other siblings* | 77,940 | - | - | - | - | - | 0.98 (0.93-1.04) | 1.000 | - | - |
| ▪ *Last son* | 8,379 | - | - | - | - | - | ref. | ref. | - | - |

**Table A3:**
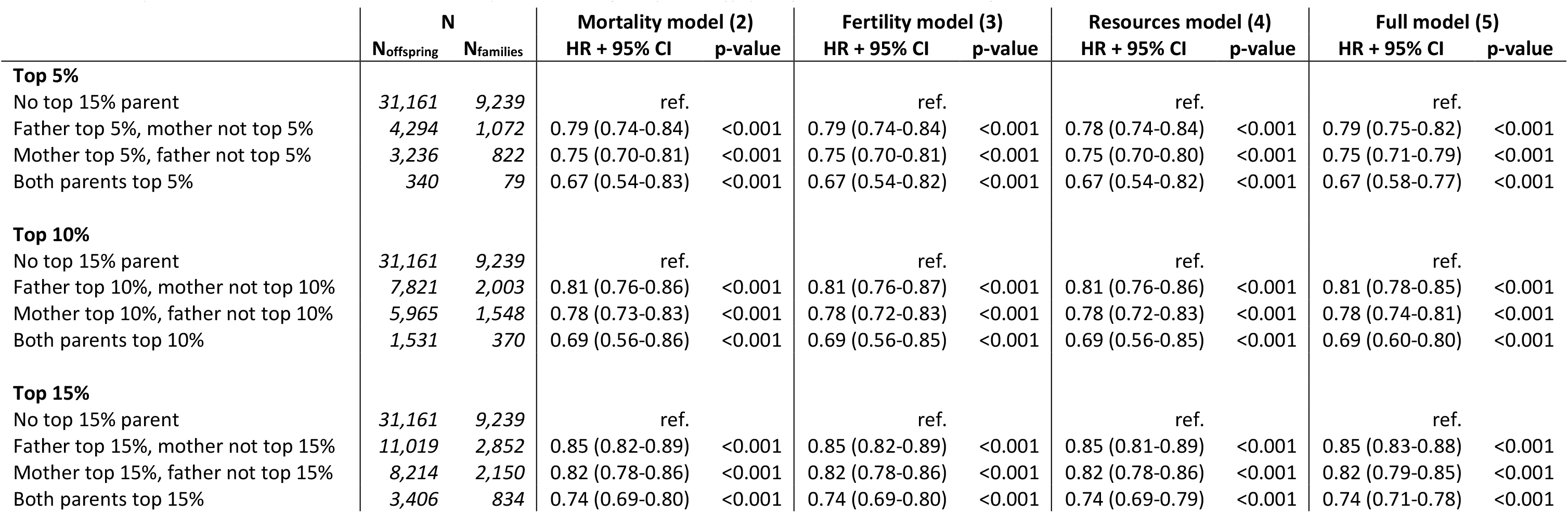
Separate model associations between parental longevity and offspring survival between ages 5-100, robustness checks

|  | N |  | Mortality model (2) |  | Fertility model (3) |  | Resources model (4) |  | Full model (5) |  |
| --- | --- | --- | --- | --- | --- | --- | --- | --- | --- | --- |
|  | N<sub>offspring</sub> | N<sub>families</sub> | HR + 95% CI | p-value | HR + 95% CI | p-value | HR + 95% CI | p-value | HR + 95% CI | p-value |
| **Top 5%** |  |  |  |  |  |  |  |  |  |  |
| No top 15% parent | 31,161 | 9,239 | ref. |  | ref. |  | ref. |  | ref. |  |
| Father top 5%, mother not top 5% | 4,294 | 1,072 | 0.79 (0.74-0.84) | <0.001 | 0.79 (0.74-0.84) | <0.001 | 0.78 (0.74-0.84) | <0.001 | 0.79 (0.75-0.82) | <0.001 |
| Mother top 5%, father not top 5% | 3,236 | 822 | 0.75 (0.70-0.81) | <0.001 | 0.75 (0.70-0.81) | <0.001 | 0.75 (0.70-0.80) | <0.001 | 0.75 (0.71-0.79) | <0.001 |
| Both parents top 5% | 340 | 79 | 0.67 (0.54-0.83) | <0.001 | 0.67 (0.54-0.82) | <0.001 | 0.67 (0.54-0.82) | <0.001 | 0.67 (0.58-0.77) | <0.001 |
| **Top 10%** |  |  |  |  |  |  |  |  |  |  |
| No top 15% parent | 31,161 | 9,239 | ref. |  | ref. |  | ref. |  | ref. |  |
| Father top 10%, mother not top 10% | 7,821 | 2,003 | 0.81 (0.76-0.86) | <0.001 | 0.81 (0.76-0.87) | <0.001 | 0.81 (0.76-0.86) | <0.001 | 0.81 (0.78-0.85) | <0.001 |
| Mother top 10%, father not top 10% | 5,965 | 1,548 | 0.78 (0.73-0.83) | <0.001 | 0.78 (0.72-0.83) | <0.001 | 0.78 (0.72-0.83) | <0.001 | 0.78 (0.74-0.81) | <0.001 |
| Both parents top 10% | 1,531 | 370 | 0.69 (0.56-0.86) | <0.001 | 0.69 (0.56-0.85) | <0.001 | 0.69 (0.56-0.85) | <0.001 | 0.69 (0.60-0.80) | <0.001 |
| **Top 15%** |  |  |  |  |  |  |  |  |  |  |
| No top 15% parent | 31,161 | 9,239 | ref. |  | ref. |  | ref. |  | ref. |  |
| Father top 15%, mother not top 15% | 11,019 | 2,852 | 0.85 (0.82-0.89) | <0.001 | 0.85 (0.82-0.89) | <0.001 | 0.85 (0.81-0.89) | <0.001 | 0.85 (0.83-0.88) | <0.001 |
| Mother top 15%, father not top 15% | 8,214 | 2,150 | 0.82 (0.78-0.86) | <0.001 | 0.82 (0.78-0.86) | <0.001 | 0.82 (0.78-0.86) | <0.001 | 0.82 (0.79-0.85) | <0.001 |
| Both parents top 15% | 3,406 | 834 | 0.74 (0.69-0.80) | <0.001 | 0.74 (0.69-0.80) | <0.001 | 0.74 (0.69-0.79) | <0.001 | 0.74 (0.71-0.78) | <0.001 |

**Table A4:** Separate model associations between parental longevity and offspring survival between ages 0-5, robustness checks

|  | N |  | Mortality model (2) |  | Fertility model (3) |  | Resources model (4) |  | Full model (6) |  |
| --- | --- | --- | --- | --- | --- | --- | --- | --- | --- | --- |
|  | N<sub>offspring</sub> | N<sub>families</sub> | HR + 95% CI | p-value | HR + 95% CI | p-value | HR + 95% CI | p-value | HR + 95% CI | p-value |
| **Top 5%** |  |  |  |  |  |  |  |  |  |  |
| No top 5% parent | 87,780 | 14,766 | ref. |  | ref. |  | ref. |  | ref. |  |
| Father top 5%, mother not top 5% | 7,782 | 1,166 | 0.87 (0.82-0.93) | <0.001 | 0.88 (0.82-0.94) | <0.001 | 0.87 (0.81-0.94) | <0.001 | 0.89 (0.85-0.93) | <0.001 |
| Mother top 5%, father not top 5% | 5,473 | 881 | 0.80 (0.74-0.86) | <0.001 | 0.77 (0.71-0.84) | <0.001 | 0.75 (0.69-0.81) | <0.001 | 0.80 (0.76-0.84) | <0.001 |
| Both parents top 5% | 542 | 82 | 0.71 (0.56-0.91) | <0.001 | 0.71 (0.55-0.92) | 0.002 | 0.67 (0.51-0.87) | <0.001 | 0.73 (0.62-0.86) | <0.001 |
| **Top 10%** |  |  |  |  |  |  |  |  |  |  |
| No top 10% parent | 74,336 | 12,639 | ref. |  | ref. |  | ref. |  | ref. |  |
| Father top 10%, mother not top 10% | 14,381 | 2,207 | 0.89 (0.84-0.95) | <0.001 | 0.90 (0.84-0.96) | <0.001 | 0.89 (0.83-0.96) | <0.001 | 0.91 (0.87-0.95) | <0.001 |
| Mother top 10%, father not top 10% | 10,383 | 1,665 | 0.82 (0.75-0.88) | <0.001 | 0.79 (0.73-0.86) | <0.001 | 0.77 (0.70-0.83) | <0.001 | 0.82 (0.77-0.86) | <0.001 |
| Both parents top 10% | 2,477 | 394 | 0.73 (0.57-0.93) | 0.003 | 0.73 (0.57-0.95) | 0.006 | 0.68 (0.52-0.90) | <0.001 | 0.74 (0.63-0.88) | <0.001 |
| **Top 15%** |  |  |  |  |  |  |  |  |  |  |
| No top 15% parent | 60,963 | 10,538 | ref. |  | ref. |  | ref. |  | ref. |  |
| Father top 15%, mother not top 15% | 20,388 | 3,155 | 0.91 (0.87-0.95) | <0.001 | 0.91 (0.87-0.95) | <0.001 | 0.91 (0.87-0.96) | <0.001 | 0.92 (0.89-0.94) | <0.001 |
| Mother top 15%, father not top 15% | 14,511 | 2,314 | 0.85 (0.81-0.89) | <0.001 | 0.83 (0.79-0.88) | <0.001 | 0.81 (0.77-0.86) | <0.001 | 0.85 (0.82-0.88) | <0.001 |
| Both parents top 15% | 5,715 | 898 | 0.78 (0.72-0.84) | <0.001 | 0.76 (0.70-0.82) | <0.001 | 0.75 (0.68-0.81) | <0.001 | 0.78 (0.74-0.82) | <0.001 |

**Table A5:** Full model (5) Association between having a top 10% parent and offspring survival for offspring who did not lose a parent before age 5, between ages 5-100

|  | Sample, no selections |  |  |  | Both parents alive at age 5 |  |  |  |
| --- | --- | --- | --- | --- | --- | --- | --- | --- |
|  | N |  | Full model (5) |  | N |  | Full model (5) |  |
|  | N<sub>offspring</sub> | N<sub>families</sub> | HR + 95% CI | p-value | N<sub>offspring</sub> | N<sub>families</sub> | HR + 95% CI | p-value |
| **Parental longevity** |  |  |  |  |  |  |  |  |
| No top 5% parent | 38,483 | 11,154 | ref. | ref. | 34,279 | 6,499 | ref. | ref. |
| Father top 5% | 7,821 | 2,003 | 0.83 (0.81-0.86) | <0.001 | 7,437 | 1,225 | 0.83 (0.81-0.86) | <0.001 |
| Mother top 5% | 5,965 | 1,548 | 0.80 (0.77-0.83) | <0.001 | 5,582 | 1,002 | 0.81 (0.78-0.84) | <0.001 |
| Both parents top 5% | 1,531 | 370 | 0.75 (0.71-0.81) | <0.001 | 1,531 | 247 | 0.76 (0.71-0.81) | <0.001 |

HR = hazard ratio, CI = confidence interval. Reported HRs result from the full model (5) and confidence intervals and p-values are corrected for the number of variables in the dataset.
Full model includes parental longevity, sex, and birth cohort.

**Table A6:** Full model (6) associations between having a top 10% parent and offspring survival for offspring who did not lose a parent before age 5, between ages 0-5

|  | Sample, no selections |  |  |  | Both parents alive at age 5 |  |  |  |
| --- | --- | --- | --- | --- | --- | --- | --- | --- |
|  | N |  | Full model (5) |  | N |  | Full model (5) |  |
|  | N<sub>offspring</sub> | N<sub>families</sub> | HR + 95% CI | p-value | N<sub>offspring</sub> | N<sub>families</sub> | HR + 95% CI | p-value |
| **Parental longevity** |  |  |  |  |  |  |  |  |
| No top 5% parent | 74,336 | 12,639 | ref. | ref. | 64,172 | 11,215 | ref. | ref. |
| Father top 5% | 14,381 | 2,207 | 0.93 (0.90-0.96) | <0.001 | 13,434 | 2,066 | 0.94 (0.91-0.97) | <0.001 |
| Mother top 5% | 10,383 | 1,665 | 0.85 (0.82-0.89) | <0.001 | 9,627 | 1,581 | 0.89 (0.85-0.92) | <0.001 |
| Both parents top 5% | 2,477 | 394 | 0.78 (0.72-0.84) | <0.001 | 2,477 | 394 | 0.82 (0.76-0.88) | <0.001 |

HR = hazard ratio, CI = confidence interval. Reported HRs result from the full model (5) and confidence intervals and p-values are corrected for the number of variables in the dataset.
Full model includes parental longevity, sex, birth cohort, sibling infant mortality, descending from a smalls sibship, parental birth spacing, age father at birth, age mother at birth, paternal socioeconomic status, and number of siblings alive at birth.

**Table A7:** Association between birth spacing and offspring survival for small (2-3) and large (9+) sibships between ages 0-5 in the full model (5)

|  | 2-3 siblings |  |  |  | 9+ siblings |  |  |  |
| --- | --- | --- | --- | --- | --- | --- | --- | --- |
|  | N |  | Full model (6) |  | N |  | Full model (6) |  |
|  | N<sub>offspring</sub> | N<sub>families</sub> | HR + 95% CI | p-value | N<sub>offspring</sub> | N<sub>families</sub> | HR + 95% CI | p-value |
| **Parental birth spacing** |  |  |  |  |  |  |  |  |
| <1.5 | 1,933 | 798 | ref. | ref. | 13,694 | 1,087 | ref. | ref. |
| 1.5-2.5 | 3,000 | 1,202 | 0.87 (0.80-0.94) | <0.001 | 31,938 | 2,959 | 0.81 (0.79-0.83) | <0.001 |
| >2.5 | 3,273 | 1,301 | 0.69 (0.64-0.75) | <0.001 | 1,769 | 191 | 0.72 (0.69-0.75) | <0.001 |

HR = hazard ratio, CI = confidence interval. Reported HRs result from the full model (5) and confidence intervals and p-values are corrected for the number of variables in the dataset.
Full model includes parental longevity, sex, birth cohort, sibling infant mortality, descending from a smalls sibship, parental birth spacing, age father at birth, age mother at birth, paternal socioeconomic status, and number of siblings alive at birth.

